# Testing the Role of Δ^9^-Tetrahydrocannabinol During Adolescence as a Gateway Drug: Behavioural, Brain Imaging and Transcriptomic Studies

**DOI:** 10.1101/2020.10.19.345322

**Authors:** Javier Orihuel, Roberto Capellán, David Roura-Martínez, Marcos Ucha, Laura Gómez-Rubio, Claudia Valverde, Marta Casquero-Veiga, María Luisa Soto-Montenegro, Manuel Desco, Marta Oteo Vives, Marta Ibáñez Moragues, Natalia Magro Calvo, Miguel Ángel Morcillo, Emilio Ambrosio, Alejandro Higuera-Matas

## Abstract

Cannabis is widely consumed by adolescents, and is also a potential prior step leading to the use of other drugs later in life (Gateway Hypothesis); however, the evidence for this hypothesis is controversial. This work aimed to increase our understanding of the long-term consequences of adolescent exposure to Δ^9^-tetrahydrocannabinol (THC) and to test the Gateway Hypothesis, experimentally. We exposed rats of both sexes to THC and studied its effects on reward-related processes, brain morphology (MRI), metabolism (^1^H-MRS), function (PET) and the transcriptomic profiles of the nucleus accumbens (RNASeq). Lastly, we studied cocaine-induced cellular activation (c-Fos) and cocaine addiction-like behaviours. THC exposure increased Pavlovian to instrumental transfer in males, goal-tracking (regardless of the sex) and impulsivity, but did not affect habit formation. Adolescent THC reduced striatal volume (in females), commissural integrity and ventricular volume. Also, there were lower levels of choline compounds in the cortex of THC-exposed rats and cerebellar hypoactivation in THC-females. THC also modified some of the gene expression programs of the nucleus accumbens, which could contribute to the behavioural features observed. Lastly, THC exposure increased cocaine-induced c-Fos levels in cortical and hypothalamic areas and increased the motivation for cocaine, followed by a higher rebound of use in THC-females after reestablishing low-effort conditions. Critically, acquisition of cocaine self-administration, compulsive seeking, intake under extended access or the incubation of seeking were unaltered. These results suggest that adolescent THC exposure alters psychological and brain development and that the Gateway Hypothesis does not entirely pass the test of preclinical enquiry.

## INTRODUCTION

Adolescence is a crucial period of development, characterised by profound changes in psychological and neural processes (1–3). Consequently, the use and abuse of cannabinoids (the drugs -other than alcohol and tobacco-most widely consumed by adolescents(4)) during this period may have long-lasting consequences at the psychological and neurobiological level (5–8). Given the actual tendency of several administrations to move towards the legalisation of cannabis, it is of the utmost importance to increase our scientific understanding of the effects of this drug in the developing adolescent individual (5). A particularly concerning consequence of adolescent cannabis consumption is the potential of the drug to act as a gateway substance, facilitating the use of or addiction to other drugs later in life (termed Gateway Hypothesis, which is under intense debate (9–17)). Previous experiments by our group and others have suggested that animals with adolescent cannabinoid exposure show increased morphine (18), heroin (19, 20), fentanyl (but not oxycodone) (21) and cocaine self-administration [19, 20 but see 21], with an interesting sex-dependence of some these results. In spite of these exciting initial findings, these aforementioned studies have not always examined the complex full array of behaviours that are indicative of addiction, especially compulsive seeking or taking (25, 26). This is, arguably, a critical prerequisite to provide an appropriate answer to the debate about early cannabis use and, subsequently, drug addiction (not just drug use). In addition, the development of addiction is conditioned by several psychological processes that could also be affected by cannabis use during adolescence. These processes include (but are not limited to) pavlovian to instrumental transfer (i.e. the ability of classically conditioned cues to affect instrumental responses; PIT) (27), pavlovian conditioned approach (a measure of incentive salience; PCA) (28) and habit formation (29, 30). Also, impulsivity is a core endophenotype that predicts the development of (cocaine) addiction (31, 32).

The neural basis of this increased susceptibility to drug reward after adolescent cannabinoid exposure is poorly understood. Some previous reports, including our own, have adopted a targeted approach, focusing on the opioid (19, 33–35), endocannabinoid (35–37), glutamatergic and GABAergic (36) and dopaminergic (38) systems, but a more comprehensive study of gene networks altered by adolescent cannabinoid exposure is only recently being investigated. However, of the two studies that have adopted a transcriptome-wide sequencing approach, one did not study the nucleus accumbens (39), an essential locus of cocaine-induced rewards, and the other used a synthetic cannabinoid (WIN 55,512-2) and not the actual psychoactive component of cannabis (40).

Moreover, in addition to gaining a fuller understanding of the neurochemical alterations that may follow the adolescent consumption of cannabis, it is essential to determine if there are structural or functional alterations in the brain as a whole, in the in vivo situation. The use of neuroimaging techniques is especially useful for this purpose. While several reports in the human literature have documented long-lasting effects of cannabis consumption on brain structure and function during adolescence (41, 42), there is a paucity of this type of research using animal models. In two previous reports by our group, we used positron emission tomography -PET-imaging to study the long-term functional consequences of adolescent cannabinoid exposure on adult brain function finding alterations in the frontal and amygdalo-entorhinal cortices, the septal nuclei and the striatum (23, 43). However, no structural imaging or metabolic studies have been performed up to this date.

Considering all this, in this work, we have adopted a multidisciplinary perspective combining different neuroimaging techniques (magnetic resonance imaging -MRI-, diffusion tensor imaging -DTI, proton magnetic resonance spectroscopy -^1^H-MRS- and PET) along with a comprehensive behavioural characterisation (a study of PIT, PCA, habit formation and impulsivity) and a transcriptomic analysis (RNASeq) of the nucleus accumbens, together with studies of cocaine-induced cell activation (c-Fos) and a multiparametric cocaine self-administration protocol designed to measure several core features of addiction, to subject the Gateway Hypothesis to an extensive experimental examination.

## MATERIALS AND METHODS

A fully detailed account of the materials and methods used are provided in the Supplementary Online Material -SOM-.

### 1. Animals and THC treatment

Wistar albino rats from Charles-River S.A. (Saint-Germain-sur-l’Arbresle, France) were mated and their male and female offspring were used. All animals were maintained at a constant temperature (20±2°C) under a reverse 12-h/12-h dark/light cycle (lights on at 20:00 h), with free access to water and food (commercial diet for rodents SAFE, Augy, France), unless otherwise specified. Animals from 35 litters were assigned to different experiments, thus minimising litter effects.

Rats received nine intraperitoneal injections (2 mL/kg) of Δ^9^-THC at a dose of 3 mg/kg or vehicle (2 mL/kg) beginning in early adolescence every other day from PND 28 to PND 44 (or from PND 38 to PND 54 in the PET experiment). We used separate batches of animals for each of the experiments described below.

### 2. Pavlovian conditioned approach and habit formation

At approximately PND 90, rats were food-restricted and their weight kept between 90-95% of the original in the free-feeding state and then, the PCA protocol began. Each session consisted of 25 trials in which the feeder dispensed pellets into the magazine under a variable interval 60 s schedule of reinforcement. A lever on one of the sides of the magazine was extended 8 s before the reward. This lever was retracted in the moment of reward presentation and acted as the conditioned stimulus. The other lever was present during the whole session and served as a measure of general locomotor activity. None of the levers had programmed contingencies. We used a PCA index where an animal with a value higher than 0.5 was classified as goal-tracker, whereas animals with a PCA lower than -0.5 were categorised as sign-trackers (see SOM for further details).

Ten days after final PCA session animals, we began the sensory-specific devaluation protocol to study habit formation. The rats performed a brief and extended training scheme (see SOM for further details). Every session was implemented with a single lever protracted inside the operant box. The brief training consisted of 5 consecutive daily sessions: one FR1 session, two variable-interval (VI) 30 s sessions and two VI 60 s sessions. After the training sessions, we subjected the animals to two counterbalanced, sensory-specific satiety-based, devaluation tests. In these tests, the rats were allowed to eat pellets (devalued condition) or chow food (non-devalued condition) freely for one hour before undergoing an extinction test (5 minutes). On the day after the first test, the animals were retrained under a regular VI60 session and, on the next day, another devaluation session was carried out with the other type of food (pellets or regular chow). We also performed an omission test and an extended training protocol (see SOM for further details). The devaluation index was the percentage of lever presses in the devalued condition over the lever presses in the devalued and nondevalued conditions.

### 3. Pavlovian to Instrumental Transfer and 2-Choice Serial Reaction Time Task

At around PND90, animals were food-restricted, and their weight kept between 90-95% of the original in the free-feeding state.

The PIT protocol started two days later consisting in four consecutive phases:

(1) Pavlovian training: 10 sessions of 4 cycles composed of a) 2-minute trials under a variable interval of 30 seconds (VI30) where they were exposed to a reward-associated conditioned stimulus or CS^+^ (clicker or tone, counterbalance); b) 1-minute trials without conditioned stimulus (Inter Stimulus Interval or ISI) and c) a 2-minute trial where the animals were exposed to a non-rewarded stimulus or CS^-^ (clicker or tone, counterbalanced) (2) Instrumental training: 1 session under fixed ratio (FR) 1, three sessions under variable-ratio VR5 and three sessions under variable-ratio VR10. (3) Extinction: Two daily sessions of 20 minutes with levers protracted but no programmed consequences. (4) PIT test: One single session that began with a third extinction period of 20 minutes then 4 cycles composed CS or ISI presentations (counterbalanced) lasting two minutes. PIT was calculated as the percentage of active lever presses during the CS^+^ over the number of active lever presses during CS^+^ and CS^-^.

Ten days after the end of the PIT animals were again food-restricted and the two-choice serial reaction time task (2-CSRTT) began (following an adaptation of protocol published by Bari and Robins(44)). The primary variable measure here was the increase in premature responding in the test sessions (long inter-trial interval sessions -see SOM) over the premature responding percentage of the previous day.

### 4. Magnetic Resonance

At approximately PND 90, T2-weighted (T2-W) spin-echo anatomical images were acquired in a Bruker Pharmascan system (Bruker Medical Gmbh, Ettlingen, Germany). Volumetric analyses were made by manually segmenting the different regions of interest (ROIs) in each image by a blind experimenter and then calculating the area with ImageJ. The total brain volume and relative volume of specific areas and ventricles were extracted. Diffusion-weighted images (DTI) were acquired in the same imaging session with a spin-echo single-shot echo-planar imaging (EPI) pulse sequence. Fractional anisotropy (FA), mean diffusivity (MD), trace, the eigenvalues and eigenvector maps were calculated with a native software application written in Matlab (R2007a). The values of these indices were extracted from maps in selected Regions of interest (ROIs) with the ImageJ software. In addition, an in vivo ^1^H magnetic resonance spectroscopy study was performed in two brain regions: cortex and striatum (see SOM).

### 5. Positron Emission Tomography

PET-CT studies were performed at PND32-33 and PND60, using an Argus PET/CT scanner (SEDECAL, Madrid, Spain). We performed these two PET scans to study if the normal developmental patterns observed in control animals would be altered by THC exposure during adolescence. Static PET imaging was obtained for 45 min at 30 min post intravenous administration with 176 ± 37 MBq/kg body weight of 2-deoxy-2-[^18^F]fluoro-D-glucose ([^18^F]FDG). PET data were reconstructed using a 2D-OSEM algorithm (16 subsets and 3 iterations) with random and scatter corrections.

Four brain ROIs (hippocampus, prefrontal cortex, caudate nucleus and cortex) were segmented. Also, voxel-based 2-sample t-tests (p<0.05 uncorrected) were performed with Statistical Parametric Mapping (SPM) software (http://www.fil.ion.ucl.ac.uk/spm/software/spm12/) for each sex.

### 6. Transcriptomic analysis of the nucleus accumbens (shell)

After adolescent THC/ VEH treatment, rats were sacrificed at PND 90, the NAcc shell dissected out on ice and subjected to Illumina RNA sequencing. We used the Chipster analysis suite (45) to perform data processing and analysis.

Gene ontologies and pathways enrichment and overrepresentation were calculated with Metascape (46) (http://metascape.org) for every comparison and gene subset obtained (see SOM for further details).

### 7. c-Fos immunohistochemistry

At PND 90, rats were injected i.p. either with cocaine (20 mg/kg) or saline (0.9% NaCl sterile solution; Vitulia Spain). Ninety minutes later, they were transcardially perfused, their brains extracted and cut in a cryostat, and we performed c-Fos immunohistochemistry according to standard protocols (47), with minor modifications. We used ImageJ to analyse c-Fos+ cells in a selected set of ROIs (see Table S6) (see SOM for further details).

### 8. Cocaine-addiction like behaviour

On PND 90, rats underwent a single food-reinforced FR1 instrumental training session limited to 10 reinforcers in Med-Associates operant boxes. After this, an intravenous catheter was implanted into the right jugular vein, and after recovery, the study of cocaine addiction-like behaviour began. Cocaine (0.5 mg/kg) self-administration took place in Coulburn operant boxes. The protocol consisted of 6 consecutive phases: (1) acquisition: 12 daily sessions lasting 2 hours each under an FR1 schedule; (2) motivation for consumption: 6 sessions of 2 hours under a progressive ratio (PR) schedule; (3) rebaseline: 3 sessions of 2 hours under FR1; (4) compulsive (punished) seeking: a single 1-hour session under an FR3 schedule in which the animal randomly received an infusion of cocaine or a 0.5 mA plantar shock for 0.5 s (5) extended access: ten 6-hours sessions under FR1; (6) cue-induced relapse: four 1-hour sessions with cues as in the acquisition sessions but without drug delivery, occurring after 1, 30, 60 and 100 days of forced abstinence (see SOM for further details).

### 9. Statistical Analysis

In general, for the experiments involving repeated measures, we used a mixed ANOVA with two between-subject factors (*Sex* and *Adolescent Treatment*) and one within-subject factor (*Sessions* or *Developmental Stage*). Significant interactions were followed using simple effects analysis. Mann-Witney and Kruskal-Wallis tests were used when the normality and homoscedasticity criteria for the ANOVA were not met. For the experiments without repeated measures, we used standard two-way ANOVAs (see SOM for further details).

In the PET study, we also used voxel-based 2-sample t-tests (p<0.05 uncorrected) with Statistical Parametric Mapping (SPM) software (http://www.fil.ion.ucl.ac.uk/spm/software/spm12/) for each sex. Only clusters larger than 50 adjacent voxels were considered in order to minimise the effect of type I errors.

## RESULTS

### 1. Reward-related psychological alterations induced by THC exposure during adolescence

THC-exposed rats (irrespective of the sex) were more biased to display goal-tracking behaviour than their VEH-treated controls (Figures 1A, S2 and Table S2). This notwithstanding, conditioned cues were more effective in potentiating reward seeking by instrumental responses (pavlovian-to-instrumental transfer) in THC-animals, although this effect only happened in the males (Figure 1B, S3 and Table S3). Given its role in the development of addiction, we also decided to explore if habit formation could be affected by adolescent exposure to THC. We used sensory-specific devaluation to test this hypothesis and found no differences in habit formation between VEH and THC-treated rats or between males and females (see Figures 1C, S4 and Table S5). Because of its importance as a predisposing factor to several psychopathologies, we also studied motor-waiting impulsivity in the 2-CSRTT. We found that female rats exposed to THC increased their premature responding (reflecting higher impulsive behaviour) in the first long-ITI session than the VEH-controls. Also, THC-exposed rats (regardless of the sex) were more impulsive in the second long-ITI sessions (however, this effect disappeared once the rats were already accustomed to the ITI challenge, in the third long-ITI session, suggesting that these effects also interact with the novelty of the task and reflect state-like impulsivity more than a stable trait) (Figure 1D). In addition, during the baseline training sessions (phase 12 of training), THC-exposed rats performed worse (i.e. they made less correct responses) (see Table S4), although the size of this effect was small. Moreover, this lower performance was transient, and no longer evident in the last day of training and was absent during the tests. See Figure S5 for further details.

**Figure 1:**
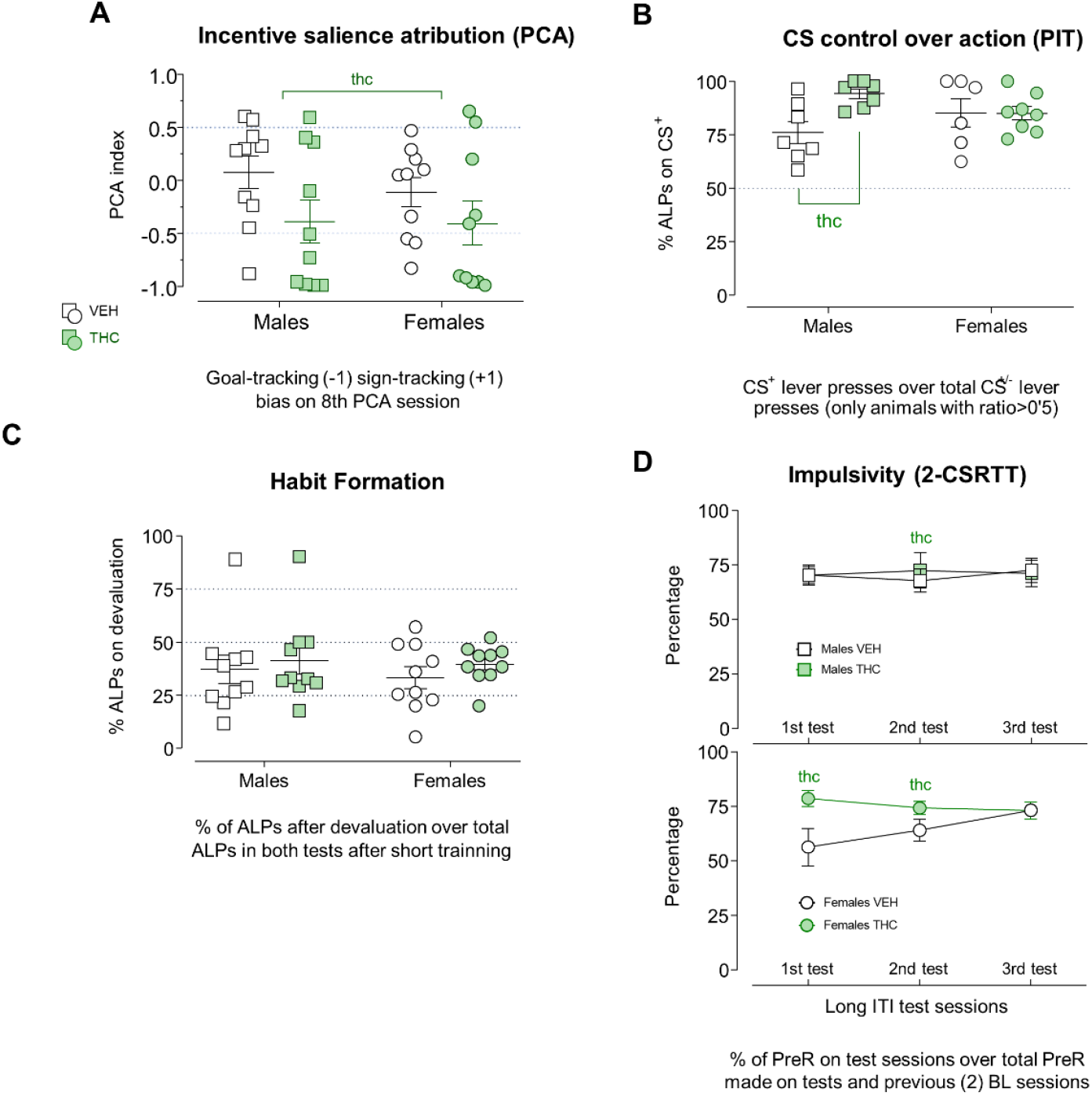
Main index measurements of the four addiction-related behavioral traits explored. Study of the effects of adolescent THC exposure on reward-related processes and impulsivity. Graphs represent individual values (dots) and mean±SEM (lines). **A)** Study of incentive salience attribution using the Pavlovian Conditioned Approach (PCA) index obtained in the 8^th^ autoshapping session. Male VEH n=10; Male THC n=10; Female VEH n=10; and Female THC n=10. Positive values indicate a bias to attribute incentive salience to outcome predictive signals, namely sign-tracking, while negative values indicate goal-tracking or the tendency to attribute salience to the goal (reward). The adolescent THC treatment biased the index towards negative values, indicating increased goal-tracking (F_1,36_=4.539; p=0.040; η_p_^2^=0.11). See Supplementary Figure 2 and Supplementary Table 2 for a more detailed account of the training procedures and the results obtained. **B)** CS^+^ LP ratio on Pavlovian to Instrumental Transfer (PIT). Male VEH n=12; Male THC n=12; Female VEH n=11; Female THC n=11. Higher percentages are obtained if instrumental responding is high during CS^+^ presentation and/or low in CS^-^ (indicating increased PIT). The Adolescent Treatment had a powerful influence of pavlovian CS control over instrumental behaviour. A general Adolescent Treatment effect was found (F_1,25_=4.685; p=0.040; η_p_^2^=0.15) suggestive of increased PIT in THC animals however, the significant Sex x Adolescent Treatment interaction (F_1,25_=4.996; p=0.035; η_p_^2^=0.16) revealed that the increase on ALPs CS^+^ ratio was only significant in THC males compared to VEH males (F_1,25_=10.108; p=0.004; η_p_^2^=0.28). Animals who did not express PIT (an index over 0.5) were excluded from this analysis. There were no differences in the acquisition of pavlovian associations or instrumental responses in the rats showing that the results obtained were specific to pavlovian motivation (see Figure S3 and Table S3 for further details concerning pavlovian and instrumental training, the PIT testing procedures and additional results). **C)** Devaluation index obtained in the outcome devaluation test after five sessions of instrumental training. Male VEH n=10; Male THC n=10; Female VEH n=10; Female THC n=10. The adolescent THC treatment had no long-term effects on this measure (F_3,36_= 0.420; p=0.740; η_p_^2^=0.03). See Figure S4 and Table S5 for a for additional results. **D)** Percentage of premature responses increase in the 2-Choice Serial Reaction Time Task (2-CSRTT) test compared to previous baseline sessions. Male VEH n=12; Male THC n=12; Female VEH n=12; Female THC n=12. The percentage of increase showed a Sex x Adolescent Treatment interaction in the first long ITI session (F_1,44_= 4.034; p= 0.051; η_p_^2^= 0.08), the simple effects analysis showed a greater increase in THC females compared to their VEH counterparts (F_1,44_= 7.892; p= 0.007; η_p_^2^= 0.15); in the second long ITI session a general Adolescent Treatment effect also obtained, indicating a more robust increase in premature responding in THC animals (F_1,44_= 5.240; p= 0.027; η_p_^2^=0.10). No factor effect was detected in the third long ITI test (F_3,44_ = 0.057; p=0.988; η_p_^2^=0.13). See Figure S4 and Table S4 for further details about the training procedures and results.

### 2. Structural and functional alterations induced by THC exposure during adolescence revealed by neuroimaging techniques

The MRI data showed that exposure to THC during adolescence leads to structural alterations evident at adulthood. More specifically, there was a reduction in the volume of the dorsal striatum observed in adult females that had a chronic THC treatment during adolescence (Figure 2). The globus pallidus was also smaller in THC-exposed animals (this effect was only significant in the right hemisphere; the effect was a trend when both hemispheres were analysed globally). Furthermore, total ventricular volume and the volume of the lateral ventricles were reduced in the THC-exposed rats (see Figure 3). In the cerebral aqueduct of Silvius, this reduced volume was only observed in THC-males (see Table S1 in the SOM for further details).

**Figure 2:**
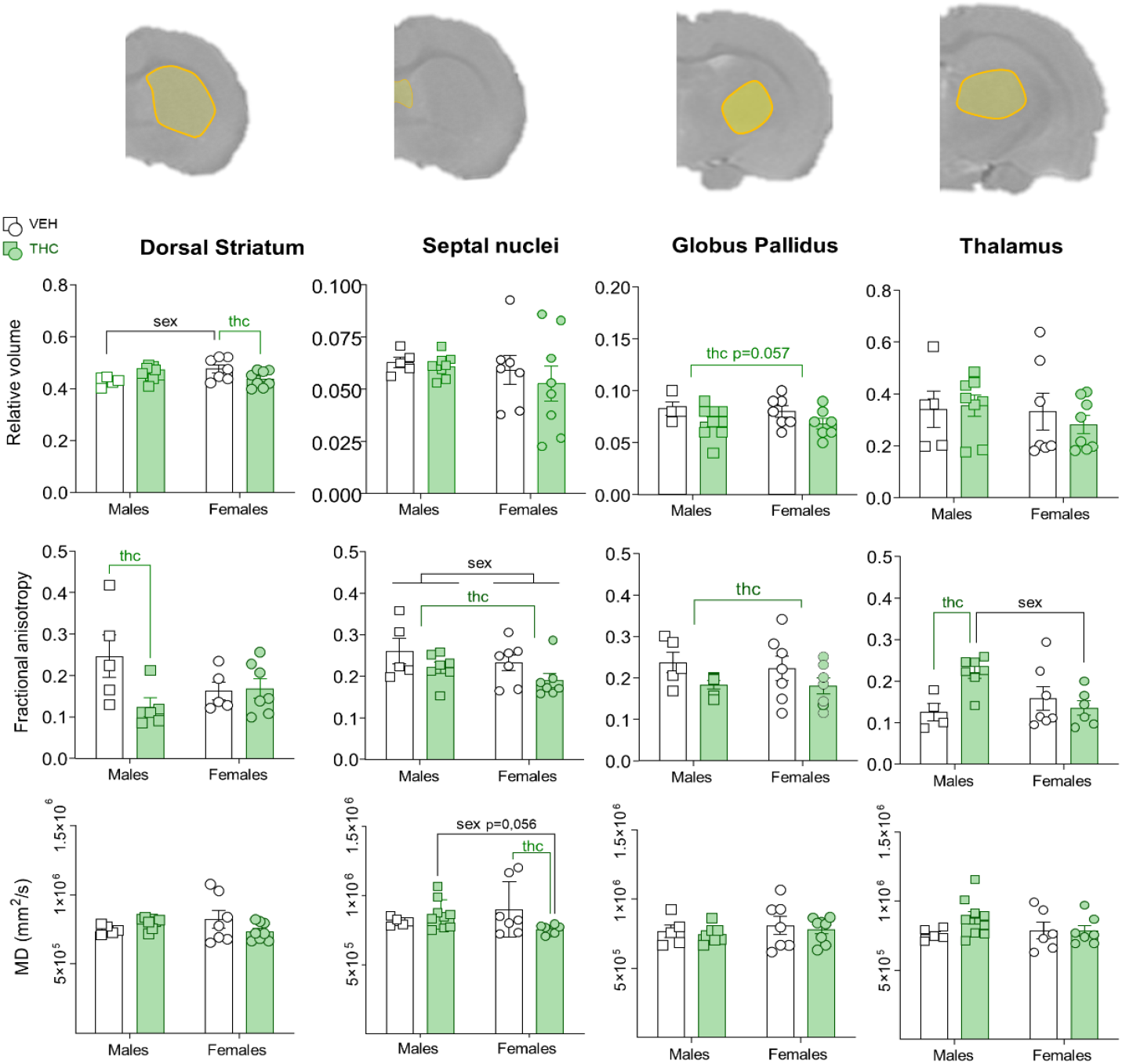
MRI Grey matter analysis. Male VEH n=5; Male THC n=9; Female VEH n=7; Female THC n=8. The most representative effects are depicted. Graphs represent individual values (dots) and Mean±SEM (bars). Within each graph, green coloured lines and “thc” represent a significant effect of the THC treatment (Adolescent Treatment). Black lines and “sex” represent statistically significant effects of the Sex factor. The columns from left to right represent the volumetric analysis, calculated as the relative volume of the structure within the sections containing it, DTI values obtained for mean diffusivity (MD) and fractional anisotropy (FA) in each of the four different structures depicted in each row; from top to bottom: Striatum (STR), Septal Nuclei (SNu), Globus Pallidus (GP) and Thalamus (THA). **First row / volumetric analysis across brain areas**. In the STR we found a significant Sex x Adolescent Treatment interaction (F_1,25_=8.806; p=0.007; η_p_^2^=0.26), with females VEH having an overall larger relative volume over males VEH (F_1,25_=5.783; p=0.024; η_p_^2^=0.19) but females THC having a decreased volume compared to females VEH (F_1,25_=7.161; p=0.013; η_p_^2^=0.22). In the GP, THC animals showed an upward trend (F_1,23_=4.022; p=0.057; η_p_^2^=0.15) that was significant only in the right GP (F_1,22_=4.494; p=0.046; η_p_^2^=0.17) (Data shown on Table S1). Other significant male>female volumetric alterations were found in left Hippocampus (HIPP), Total Cortex (Cx) and Cerebellum (CE) (see Supplementary Table 1). **Second row / FA values across brain areas**. In the anterior section of the STR, there was a Sex x Adolescent Treatment interaction (F_1,17_=6.364; p=0.022; η_p_^2^=0.27); the analysis of the simple effects showed decreased FA values in male THC rats compared to male VEH animals. We also detected another Sex x Adolescent Treatment interaction (F_1,20_=7.057; p=0.015; η_p_^2^=0.26) in the THA, this time suggesting an increased FA in THC males compared with their VH controls (F_1,20_=8.144; p=0.001; η_p_^2^=0.28) and their female counterparts (F_1,20_=8.346; p=0.009; η_p_^2^=0.29). In the caudal SNu we observed a general Sex effect (F_1,21_=4.850; p=0.039; η_p_^2^=0.18) and also decreased FA in THC rats (F_1,21_=6.999; p=0.015; η_p_^2^=2.250). Decreased FA in THC animals was also statistically significant in the GP (F_1,21_=4.309; p=0.05; η_p_^2^=0.17). **Third row / MD values across brain areas**. A Sex x Adolescent Treatment interaction was detected in rostral SNu (H=9.284; p=0.026; 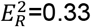), with decreased MD in THC females compared to VEH females (U=9; p=0.028; η^2^=0.34) and also lower MD in THC females compared to THC males (U=10; p=0.012; η^2^=0.39). See additional data in Table S1.

**Figure 3:**
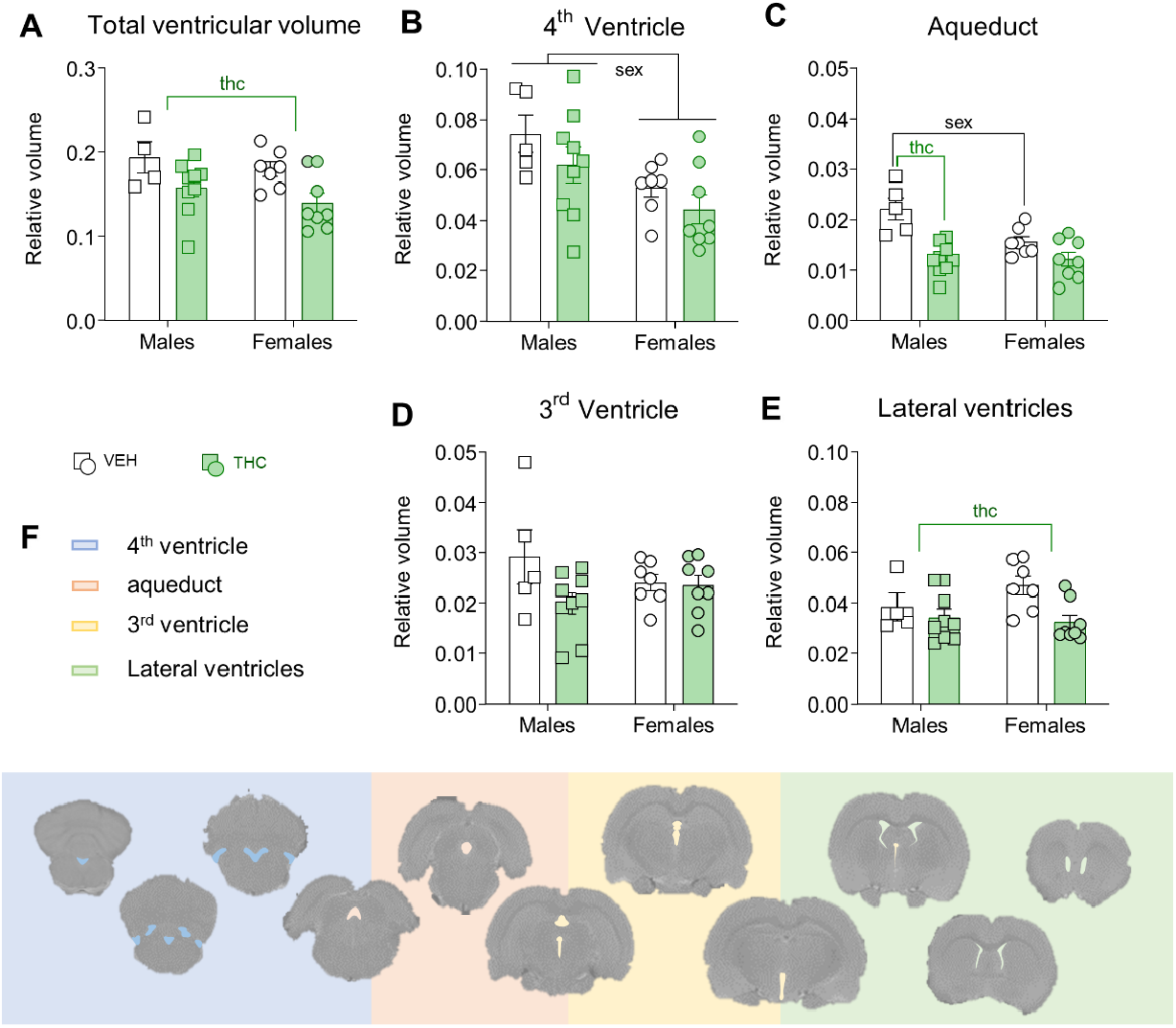
Brain Ventricle Volumetry. Graphs represent individual values (dots) and Mean±SEM (bars). **A)** The whole ventricular volume was decreased in THC-treated animals (F_1,23_=8.961; p=0.006; η_p_^2^=0.28). In order to explore where the source of this effect, we analysed the different sections of the brain ventricular system. **B)** In the fourth ventricle we only observed a male>female sexual dimorphism (F_1,25_=9.053; p=0.006; η_p_^2^=0.26). **C)** A significant Sex x Adolescent Treatment interaction (F_1,25_= 5.575; p= 0.026; η_p_^2^=0.18) surfaced in the brain aqueduct. Our follow-up analysis showed a male>female sexual dimorphism in VEH animals (F_1,25_= 9.598; p=.005; η_p_^2^=0.27), and significant differences between within the males. More specifically, THC-exposed male rats had a reduced volume (F_1,25_= 24.51; p<0.000; η_p_^2^= 0.49). **D)** In the third ventricle there was a trend for reduced volume in THC animals (F_1,25_= 3.408; p= 0.077; η_p_^2^= 0,12). **E)** In the lateral ventricles there was a reduced volume in THC animals (F_1,25_= 6.341; p= 0.019; η_p_^2^= 0.19). **F)** The different fill colours represent the ventricular area used to obtain the values represented in each corresponding graph. From caudal (left) to rostral (right): IV ventricle, aqueduct, III ventricle and lateral ventricles. The full statistical results can be examined in Supplementary Table 1.

We also analysed the integrity of defined tracks using DTI. We observed a reduced FA in the hippocampal commissure in THC-females as compared to their VEH controls. Also, THC animals (irrespective of the sex) showed decreased FA values in the rostral anterior commissure and rostral corpus callosum, possibly reflecting alterations in the integrity of these tracks (see Figure 4 and Table S1). Of note, these alterations were focused in the rostral parts of the tracks and not in the more caudal regions, where there were no significant effects.

**Figure 4:**
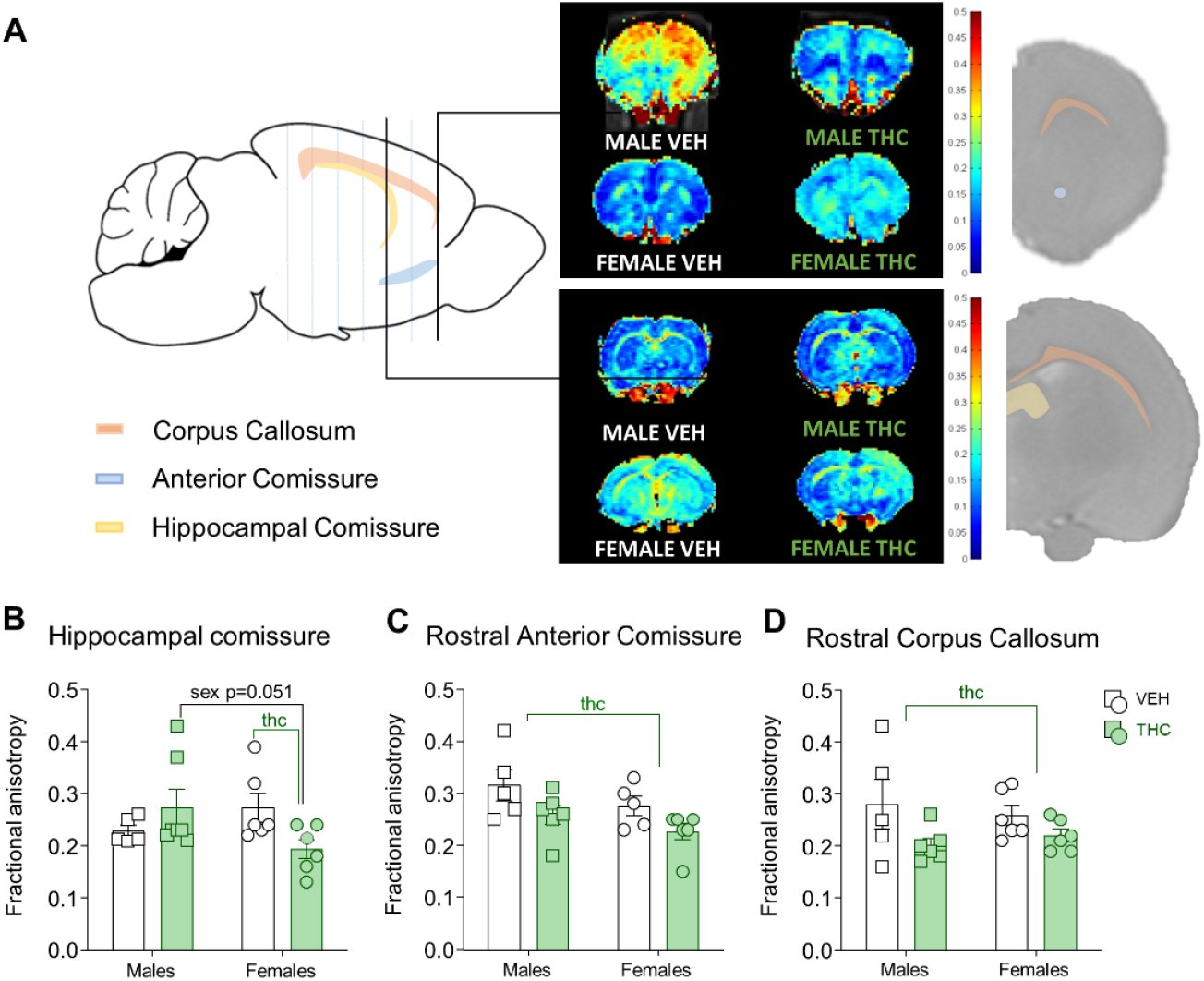
DTI FA Analysis White matter tracts. White matter Diffusion Tensor Imaging analysis of fractional anisotropy. Graphs represent individual values (dots) and Mean±SEM (bars). **A)** Representation of three major white matter tracts and the corresponding DTI FA maps where we detected significative changes in the signal. **B)** Graphs of FA values obtained in the tracts mentioned above. The FA signal in the hippocampal commissure showed a Sex x Adolescent Treatment interaction (F_1,18_= 5.537; p= 0.030; η_p_^**2**^= 0.23) which upon further analysis indicated a reduced FA in females treated with THC compared to control females (F_1,18_=5,693; p=0,028; η_p_^2^=0,240) and with their male counterparts (F_1,18_=4.359; p=0.051; η_p_^2^= 0.19). In the more rostral regions, adult animals of both sexes subjected to a chronic adolescent THC treatment had reduced FA **C)** anterior commissure (F_1,17_=5.322; p=0.034; η_p_^2^=0.23) and **D)** corpus callosum (F_1,19_= 5.298; p= 0.034; η_p_^2^= 0.23). No significant effects of the Adolescent Treatment were observed in the internal capsule (data not shown). See Table S1 for more details concerning the results of the statistical tests.

From a more functional point of view, the ^1^H-MRS data showed reduced levels of choline compounds in the cortex of THC-exposed rats (see Figure 5 and Table S1). We then turned to [^18^F]FDG PET imaging to have a more direct measure of brain function in young adult animals that were exposed to THC during adolescence. There were no statistically significant differences in SUVR values between THC and vehicle treatment in any of the regions analysed (see Table S1). We observed a developmental effect in the standardised uptake value ratio (SUVR) values that increased in all the ROIs analysed that PND 60 as compared to PND 32. We obtained a significant effect of Sex in the hippocampus and a significant Sex x Time interaction that suggested that males had higher SUVR values than females at both developmental points (see table S1). The SPM analysis of adult brain PET scans revealed some additional preliminary effects. THC-males had increased metabolism in the somatosensory cortex (S1) and the piriform cortex (Pir). In contrast, THC-exposed females showed hypometabolism in a cluster of voxels comprising the inferior colliculus and the cerebellum. There was also some marginal evidence for a hypometabolism in a cluster located at the cortex (mostly the motor and sensory cortices) (see Figure 6).

**Figure 5:**
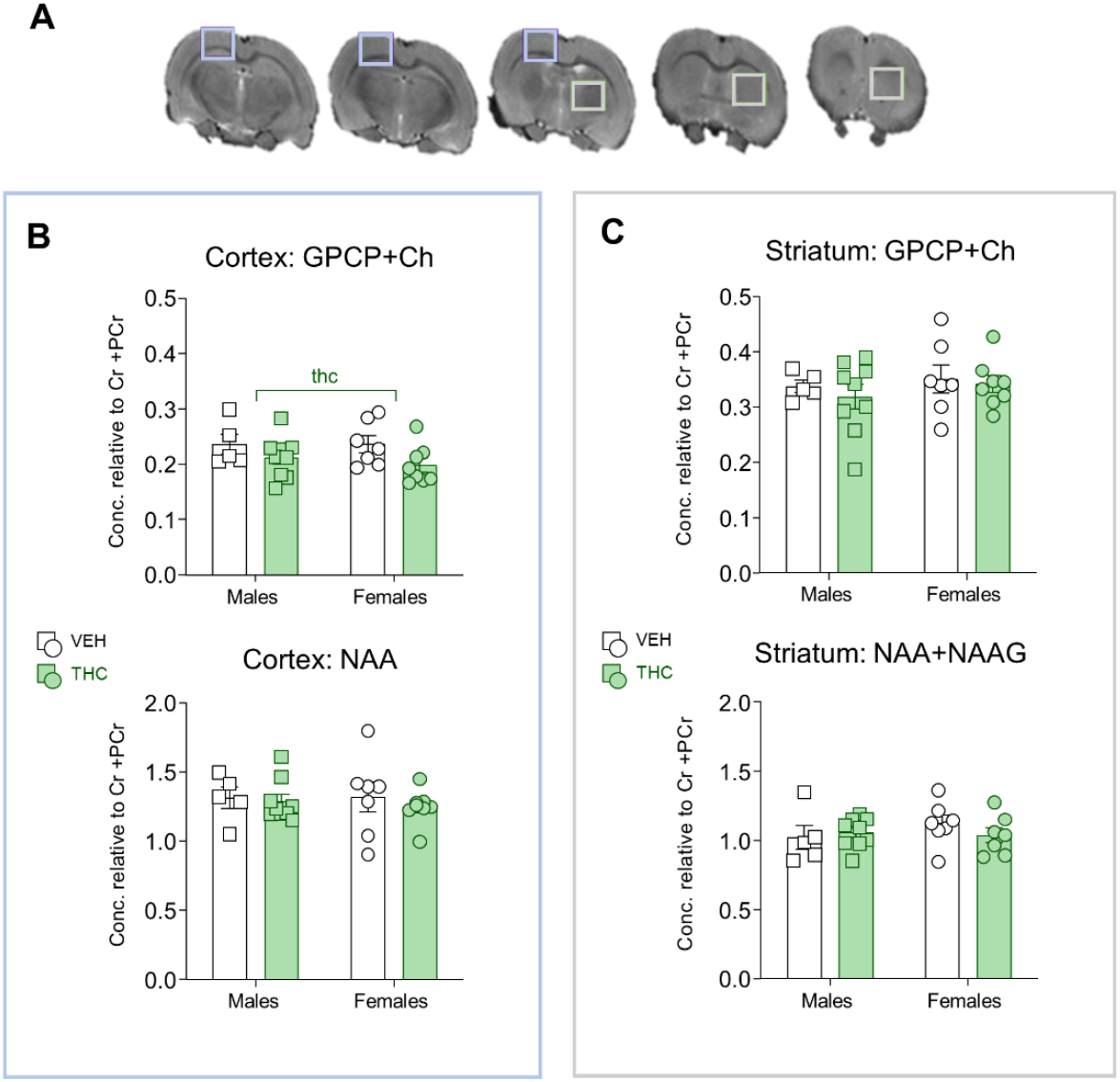
^1^H NMR Spectroscopy. The graphs represent individual values (dots) and Mean±SEM (bars). **A)** The 3 mm^3^ voxel located in the cortex (colored in blue) or the striatum (colored in grey) used to obtain the spectra. **B)** The cortical GPCP+Ch signal (glycerophosphorylcholine, phosphorylcholine, choline) was reduced in THC-treated animals (F_1,25_=4.629; p=0.041; η_p_^2^=0.15) while NAA (F_3,25_= 0.941; p= 0.436; η_p_^2^= 0.10) and NAA+NAAG values (F_3,25_= 0.298; p= 0.826; η_p_^2^= 0.03; graph not shown) remained unchanged. C) In the striatum, neither GPCP+Ch (F_3,25_=0.493; p=0.690; η_p_^2^=0.056) nor NAA (F_3,25_=0.298; p=0.826; η_p_^2^=0.03) were altered by THC treatment or showed a sex specific difference. See Table S1 for further details.

**Figure 6:**
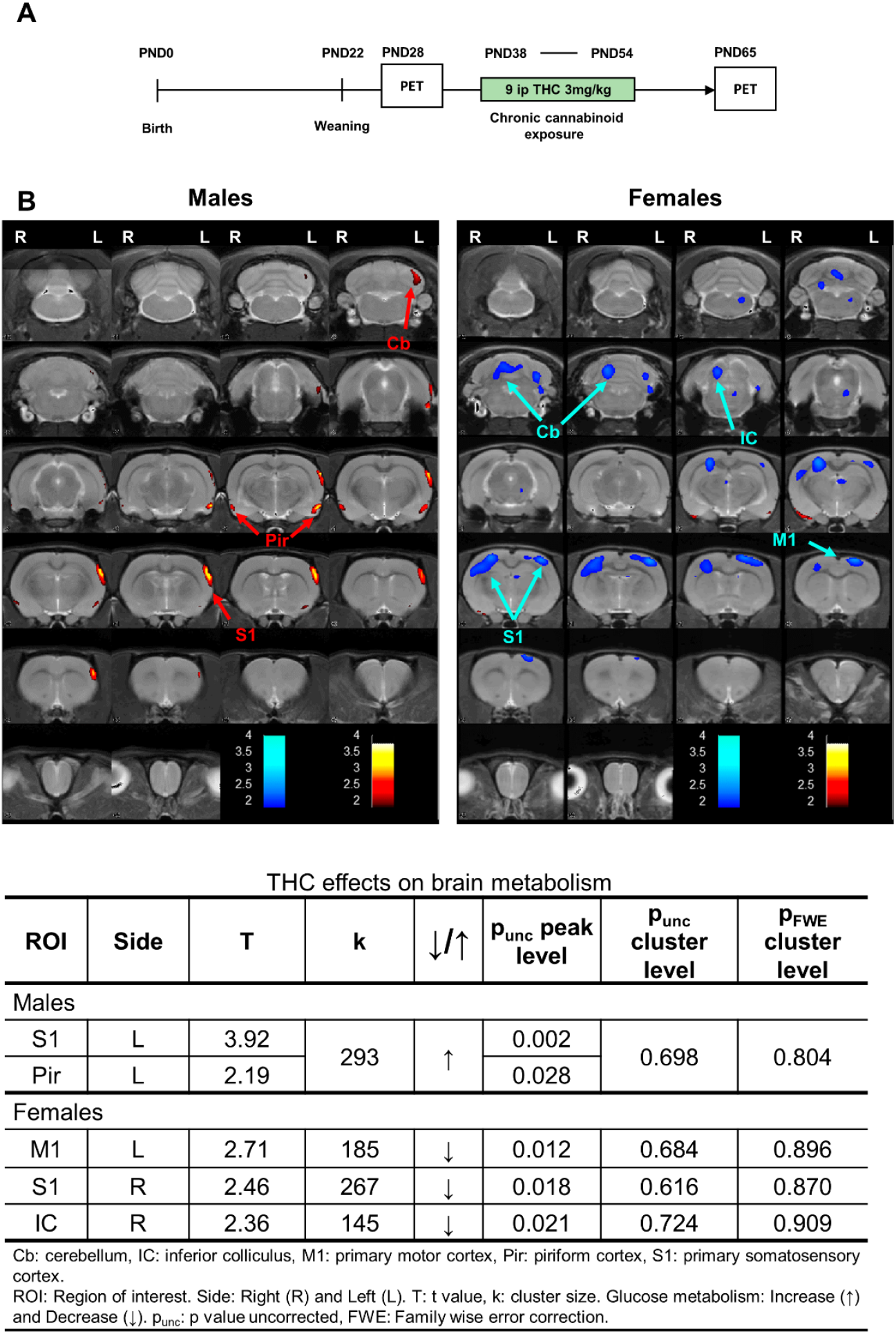
Positron emission tomography at PND65. Male VEH n=6; Male THC n=6; Female VEH n=6; Female THC n=5. THC effects on brain glucose metabolism in males and females represented as statistical parametric T-maps overlaid on a T2-image as a template. T-maps were obtained as results from 2-sample t-test analyses, applying a cluster size threshold of 50 adjacent voxels and a p-value of 0.05 (uncorrected). The colour bars represent the t-values corresponding to reductions (cold colors) or increases (hot colors) in brain metabolism. The intensity of the colour negatively correlates with the t-value of the difference in the cluster represented. The most sound effects were detected in the females where the THC treatment provoked a hypoactivation in the IC-Cb and the somatosensory Cx. ROIs: Cb, cerebellum; IC, inferior colliculus; M1, primary motor cortex; Pir, piriform cortex; S1: primary somatosensory cortex. Hemispheres: Left (L), Right (R).

### 3. THC exposure during adolescence modifies the transcriptome profile in the shell of the nucleus accumbens in a sex-specific manner

Table S8 shows the corrected p-values and the fold-change for all differentially expressed genes (DEGs). There were 96 DEGs in THC-males compared to VEH-males and 87 DEGs in the females’ comparison. Only nine of these DEGs were present in both the THC vs. VEH comparison in both sexes. Two of these nine genes (*Calb1* -Calbindin1- and *Slc17a6*-Vesicular Glutamate Transporter 2) showed higher expression in male and female rats exposed to THC. Two of them (*Dus2* - Dihydrouridine Synthase 2 -and *RGD1310819* - similar to putative protein (5S487) showed a lower expression in THC animals, and the other five showed an opposite regulation pattern in females and males (*Ttr* - Transthyretin-, *Nov* -nephroblastoma overexpressed or cellular communication network factor 3- and *Cck* -cholecystokinin-showing higher expression in THC-male rats and lower expression in the THC-females, as compared to their VEH-treated counterparts; while *Zfhx3* - zinc finger homeobox 3- and *Tenm4* - teneurin transmembrane protein 4-showed the opposite pattern).

The analysis of the gene ontologies modulated by THC exposure during adolescence reveals that, in the males, the ontology that was mainly affected was related to behavioural regulation (shown in red in Figure 7 A, B) (*Alb|Cck|Slc1a2|Scn8a|Kcnj10|Cacna1e|Nr4a2|Apba1| Calb1|Shc3|Slc12a5|Adcy1|Zfhx 3|Ttbk1|Hipk2|Crtc1*) with related ontologies such as learning and memory (*Cck|Cacna1e|Calb1|Shc3|Slc12a5|Adcy1|Ttbk1|Crtc1*), or locomotory behaviour (*Scn8a|Kcnj10|Cacna1e|Nr4a2|Apba1|Calb1|Zfhx3|Hipk2*). There is another interesting group of clusters related to the development of neuronal projections (shown in grey in Figure 7 B), including neuron projection morphogenesis (*Cck|Nr4a2|Megf8|Syt17|Plppr4|Jade2|Adcy1|Plxnb1|Bhlhe22*), and more specifically, axon development (*Cck|Nr4a2|Megf8|Plppr4|Adcy1|Plxnb1|Bhlhe22*). Other DEGs clustered around the positive regulation of neurogenesis (shown in turquoise) (*Flt1|Megf8|Syt17|Jade2|Tenm4|Plxnb1|Ttbk1|Crtc1*) and another centred around the transport of amino acids (*Cck|Slc1a2|Kcnj10|Apba1|Slc17a6*). Figures 7 C and D show the top 5 up- and down-regulated DEGs in the males (THC as compared to VEH).

**Figure 7:**
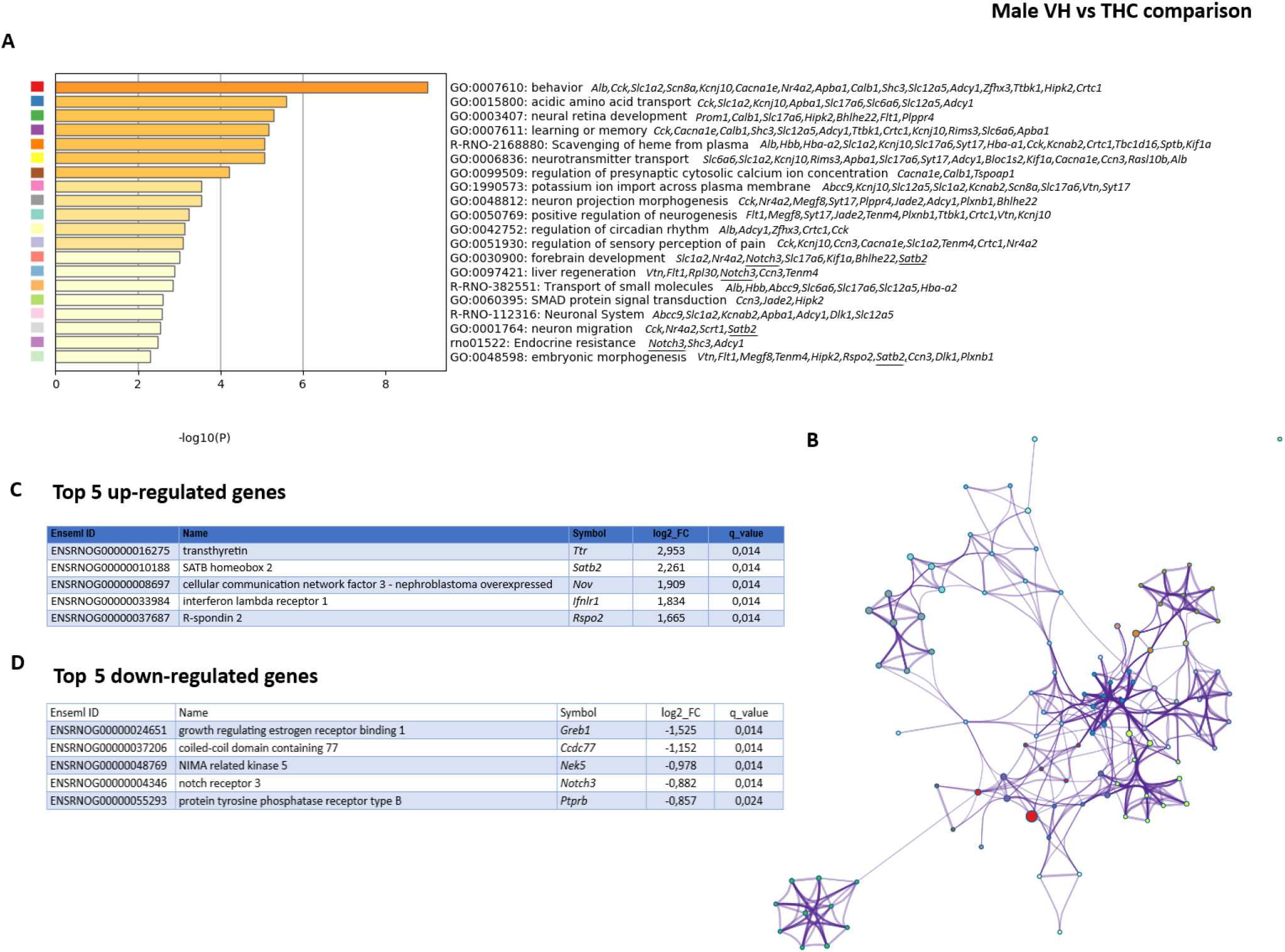
RNAseq. Main results of the NAc Shell RNASeq analysis. Male VEH n=4; Male THC n=4; Female VEH n=4; Female THC n=4. **A)** Main Gene Ontologies (or KEGG categories) affected by adolescent exposure to THC in males ranked by probability of representation. **B)** Network layout of the selected representative clusters after hierarchical grouping in the males (Figure 7) and the females (Figure 8). Each term is represented by a circle node, where its size is proportional to the number of input genes fall into that term, and its colour represents its cluster identity. Terms with a similarity score > 0.3 are linked by an edge (the thickness of the edge being proportional to the similarity score). **C)** Top five up-regulated DEGs in the males THC as compared to VEH (Figure 7) or Females THC as compared to VEH (Figure 8). Underlined genes belong to the top 5 up- or down-regulated DEGs.

In the females, there was a cluster of genes related to behavioural regulation (Figure 8A, shown in pink in Figure 8B (*Agt|Grin2d|Cck|Ptgds|Trh|Cartpt|Gal|Adcy8|Calb1|Slc17a7|Zfhx3*) that was significantly affected by adolescent THC exposure. This cluster was more restricted to the one found in males and, except for *Cck, Calb1* and *Zfhx3*, they had different components. Relatedly, another gene ontology affected in the females was the regulation of neurotransmitter levels (*Agt|Gfap|Trh|Nrn1|Slc17a6|Rgs2|Slc17a7|Sv2b|Baiap3*) and associated processes (shown in blue and purple in Figure 8B). Genes related to hormone transport and secretion were also affected (*Agt|Ttr|Trh|Cartpt|Gal|Adcy8|Ccn3|Baiap3*). Remarkably, there was an isolated constellation of gene ontologies related to microtubule reorganization (shown in red) including *Aurkb, Dnah6, Dnah1, Dnah12, Wdr63, Prc1, Ak7, Dynlrb2, Cfap44* and *Cfap43* and also with the extracellular matrix reorganization (*Agt|Gfap|Col2a1|Col5a3|Fmod*), especially collagen-related genes (*Col2a1|Col5a3|Col7a1*). Figures 8C and D show the top 5 up- and down-regulated DEGs in the females (THC as compared to VEH).

**Figure 8:**
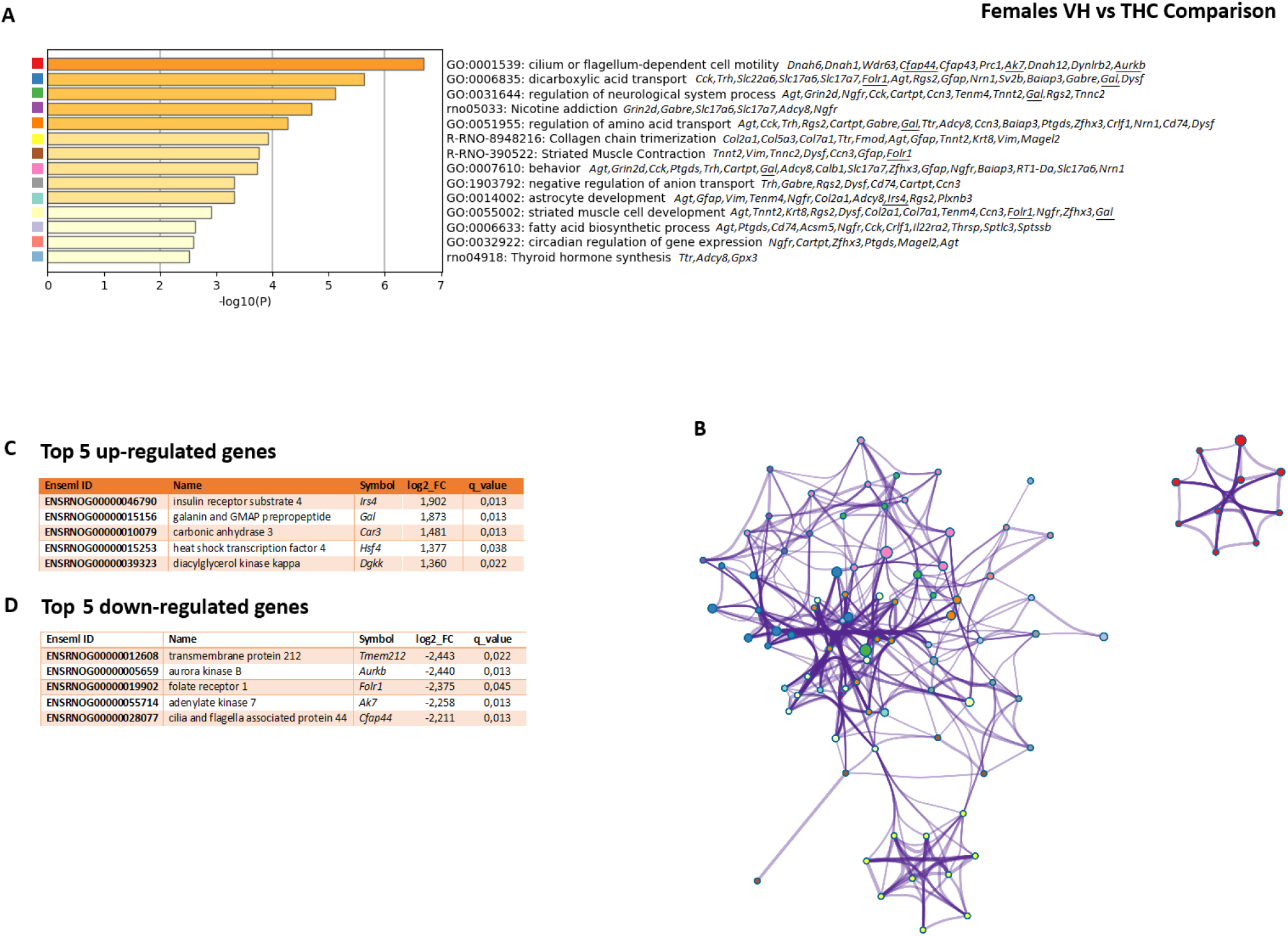
RNAseq. Main results of the NAc Shell RNASeq analysis. Male VEH n=4; Male THC n=4; Female VEH n=4; Female THC n=4. **A)** Main Gene Ontologies (or KEGG categories) affected by adolescent exposure to THC in males ranked by probability of representation. **B)** Network layout of the selected representative clusters after hierarchical grouping in the males (Figure 7) and the females (Figure 8). Each term is represented by a circle node, where its size is proportional to the number of input genes fall into that term, and its colour represents its cluster identity. Terms with a similarity score > 0.3 are linked by an edge (the thickness of the edge being proportional to the similarity score). **C)** Top five up-regulated DEGs in the males THC as compared to VEH (Figure 7) or Females THC as compared to VEH (Figure 8). Underlined genes belong to the top 5 up- or down-regulated DEGs.

### 4. Potentiated cocaine-induced c-Fos accumulation in THC-exposed rats

After studying the long-term alterations in the normal psychological and brain development after adolescent THC exposure, we then proceeded to test the Gateway Hypothesis from an experimental perspective. We first examined how the initial actions of cocaine were modified by THC exposure during adolescence. The cellular activation induced by cocaine was potentiated in the motor cortex by the THC treatment during adolescence (see Figure 9 and Table S6). In addition, we found a significant Sex x Adolescent Treatment x Adult Treatment triple interaction in the dorsomedial nucleus of the hypothalamus. The analysis of this interaction revealed a trend for cocaine to induce a significant c-Fos activity as compared to saline-injected animals in the females (but not males) exposed to THC during adolescence. There were also differences in cocaine-injected males between the THC and VEH groups. The adolescent exposure to THC also modified the mean c-Fos accumulation (irrespective of cocaine exposure) in the piriform (only in the males), retrosplenial and somatosensory cortices (see Table S6 and Figure S6).

**Figure 9:**
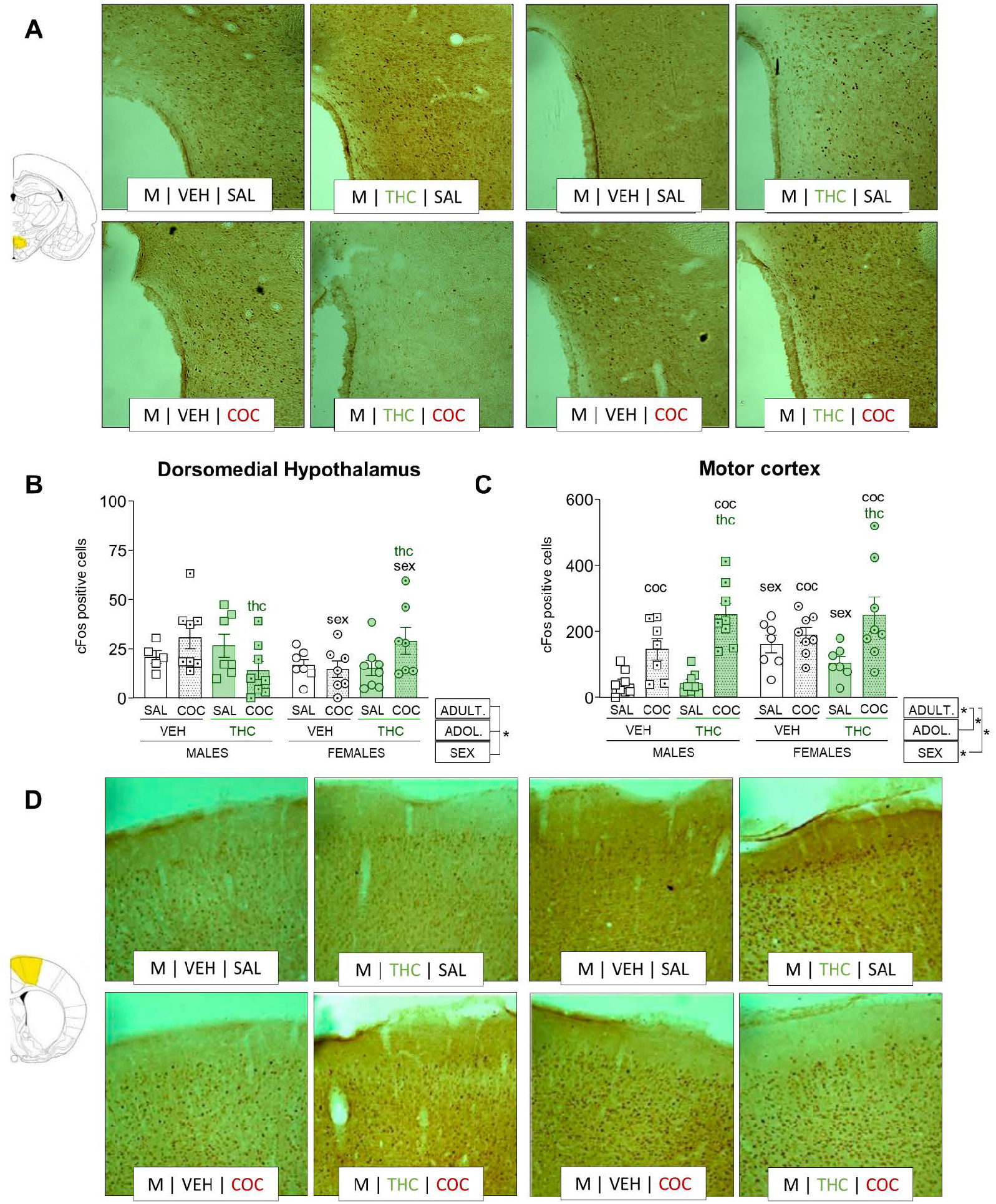
c-Fos immunohistochesmitry. c-Fos protein accumulation in response to a single i.p. cocaine (20 mg/kg) or saline injections. n=8 in all the groups. Graphs represent individual values (dots) and Mean±SEM (lines). Main factor effects are indicated with “*” next to the legend boxes on the bottom right; interactions between factors are indicated with lines joining the factors and a “*”; significant results of the analysis of the simple effect of the interactions are indicated with “sex”, “thc” or “coc” over the experimental group. *sex* stands for differences with the corresponding group of the other sex, *thc* stands for differences with the same group in the other adolescent treatment, and *coc* stands for differences with the corresponding saline group. **A)** Representative pictures of c-Fos accumulation in response to acute i.p. cocaine in the dorsomedial hypothalamic nuclei. **B)** c-Fos expression in the dorsomedial hypothalamic nuclei. We observed a significant Sex x Adolescent Treatment x Adult Treatment interaction (F_1,49_=7.148; p=0.010; η_p_^2^=0.01). Further simple effects analyses showed a trend towards significance for the effect of cocaine in THC female rats compared to their saline controls F_1,49_=3.970; p=0.055; η_p_^2^=0.08) that was absent in VEH female rats. We also found a decreased expression in cocaine-exposed VEH females compared to cocaine-exposed VEH males (F_1,49_=3.280; p=0.024; η_p_^2^=0.06). The opposite effect emerged in cocaine-exposed THC animals, with more c-Fos expression in THC females compared to THC Males (F_1,49_=3.852; p=0.036; η_p_^2^=0.07). Additionally, there was a reduced c-Fos expression in cocaine-exposed THC males compared to cocaine-exposed VEH males (F_1,49_=5.441; p=0.015; η_p_^2^=0.10). **C)** c-Fos expression in the motor cortex. There was an overall effect of Sex (F_1,53_=13.676; p=0.001; η_p_^2^=0.21) and Adult Treatment (F_1,53_=42.833; p<0.000; η_p_^2^=0.45) which indicates a higher accumulation of the c-Fos protein after cocaine. Also, there were two interactive effects, Sex x Adult Treatment (F_1,53_=7.029; p=0.011; η_p_^2^=0.12) and Adolescent Treatment x Adult Treatment (F_1,53_=5.326; p=0.025; η_p_^2^=0.09). The simple effects analysis of the Sex x Adult Treatment interaction revealed cocaine-induced c-Fos accumulation in both sexes, but based on the effect size, c-Fos induction was more robust in the males (F_1,57_=43.048; p<0.000; η_p_^2^=0.42) as compared to the females (F_1,57_=7.369; p=0.009; η_p_^2^=0.11). Confirming the main effect of Sex, we also obtained a sex-related effect in saline animals (F_1,57_=18.211; p<0.000; η_p_^2^=0.24) that was absent in cocaine-exposed rats. The simple effects analysis of the Adolescent Treatment x Adult Treatment interaction showed that, irrespective of sex, both VEH or THC animals increased cFos expression in response to cocaine. However, based on the effect sizes, response in the motor cortex is more robust in THC animals (F_1,55_=38.931; p<0.000; η_p_^2^=0.35) as compared to VEH rats (F_1,55_=9,204; p=0,004; η_p_^2^=0,12). Moreover, there was a significant difference in cocaine-exposed THC rats compared to cocaine-exposed VEH animals (F_1,57_=4.317; p=0.042; η_p_^2^=0.07). **D)** Representative pictures of c-Fos expression in response to acute i.p. cocaine in the motor cortex.

### 5. Cocaine addiction-like behaviour

Our study of cocaine addiction-like behaviours showed that the acquisition of cocaine self-administration under continuous access (fixed-ratio 1 -FR1-schedule of reinforcement) was not modified by THC exposure during adolescence. However, we observed higher cocaine intake under high-effort conditions (progressive ratio) in THC-exposed male rats (see Figure 9C, Table S7). After the progressive ratio sessions, we returned the rats to an FR1 schedule for three days. At this stage, female-THC rats showed a higher rebound in their cocaine consumption (as compared to the last acquisition sessions) (see Figure 9D, Table S7). After this, we evaluated if there was a compulsive component in the cocaine-seeking behaviour of the rats. In the punished-seeking test, all rats reduced the number of infusions achieved as compared to the last reacquisition session, but there were no effects due to Sex or Treatment (Figure 9E). After this single session, we allowed the rats to self-administer cocaine for 6 hours a day under an FR1 schedule of reinforcement for ten days. We did not observe any significant escalation in our animals (see Table S7). However, as observed in Figure 9F, the behaviour of THC-females was somewhat different from THC-males of the VEH controls. There was a surge in responding from session 5 that seemed to stabilise from session 6 to session 10. After this, we withdrew the rats from cocaine and analysed seeking responses after 1, 30, 60 and 100 days of forced withdrawal. Rats had no access to the drug during these test sessions. There were more seeking responses after 30 days than after one day of withdrawal −incubation of seeking phenomenon-as evidenced by the significant effect of Session and females showed a more robust seeking behaviour (significant effect of Sex).

**Figure 10:**
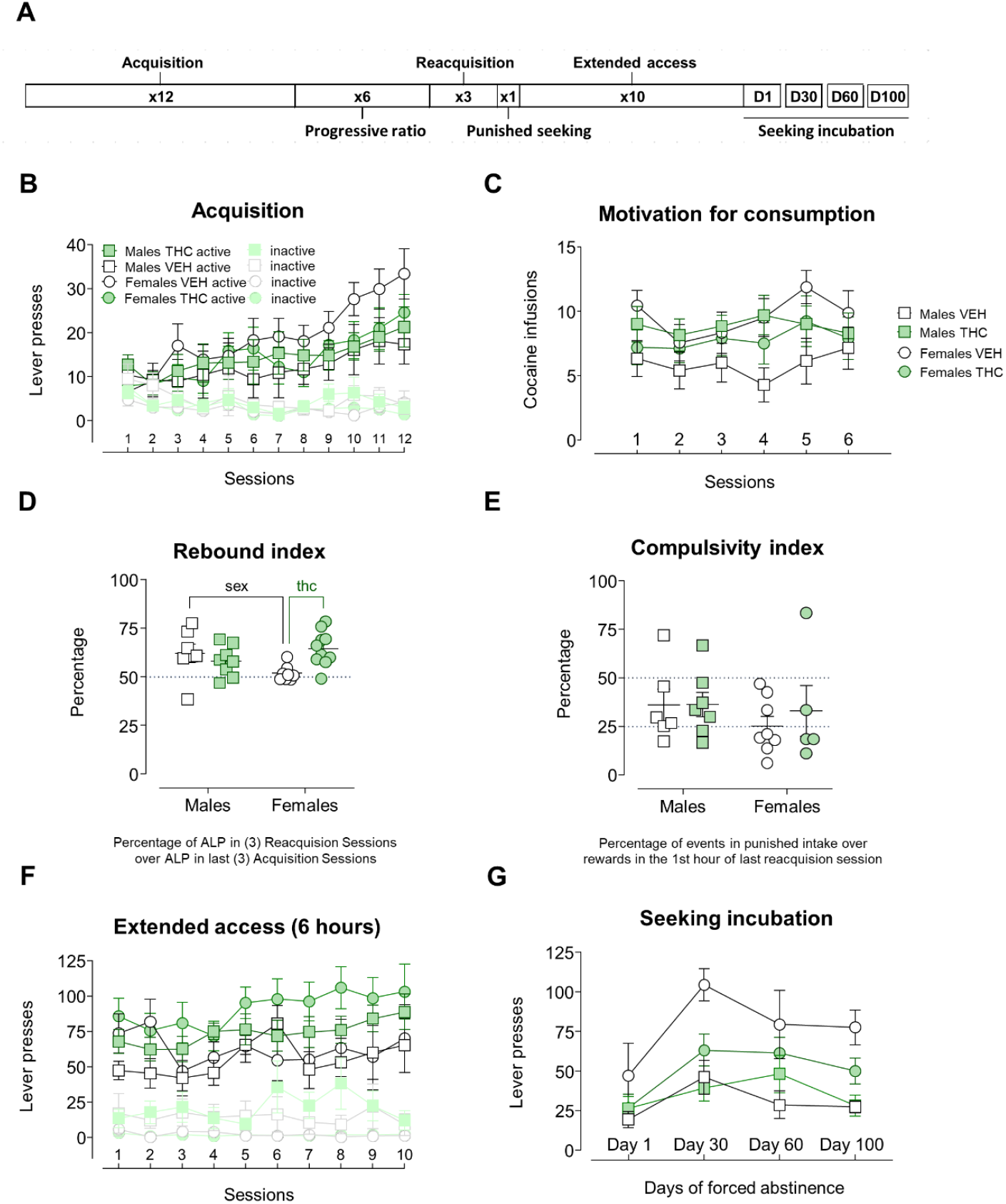
Cocaine addiction-like behaviours. Cocaine self-administration phases. Mean values are depicted with circles or squares joined by lines in the repeated measures graphs. Circles and squares represent individual values in those graphs showing a single index. Error lines express the SEM. Inital sample sizes: male VEH n=15; Male THC n=18; Female VEH n=15; Female THC n=15. **A)** Timeline of the experimental phases. **B)** Active (ALPs) and inactive lever presses (ILPs) across the twelve acquisition sessions (2h). All groups acquired a preference for the active lever (Lever: F_1,43_=39.218; p<0.000; η_p_^2^=0.48) and increased their self-administration behaviour (Lever x Session: F_4,175.3_=7.259; p<0.000; η_p_^2^=0.14) with no differences due to Sex or Adolescent Treatment. **C)** Cocaine infusions across the six progressive ratio sessions (2h). There was no Session effect in the number cocaine infusions (F_5,25_=2.021; p=0.080; η_p_^2^=0.07) but did observe a significant Sex x Adolescent Treatment interaction (F_1,25_=5.215; p=0.031; η_p_^2^=0.17). A followup analysis showed an increased number of cocaine infusions in THC males compared to VEH males (F_1,25_=6.197; p=0.032; η_p_^2^=0.38) an effect that was absent in the females. In addition, VEH females achieve a higher number of infusions compared to VEH males (F_1,25_=7.717; p=0.018; η_p_^2^=0.41). **D)** Rebound index in subsequent FR1 sessions (2h). There was a Sex x Adolescent Treatment interaction (F_1,29_=7.507; p=0.010; η_p_^2^=0.20) in the rebound of the self-administration behaviour after the high effort conditions imposed by the progressive ratio. The simple effect analysis showed that cocaine intake in VEH females remained roughly equal (around 50%) but lower compared to VEH males (62±1.8%) (F_1,29_=5.165; p=0.031; η_p_^2^=0.15), while THC females showed a higher increase in FR1 responding (as compared the last acquisition sessions) than VEH females (F_1,29_=9.497; p=0.004; η_p_^2^=0.24); however, this effect was absent in the male groups. **E)** Compulsivity index on the punished seeking phase (1h). Percentage of events achieved (shocks or cocaine infusions) normalised with the infusions achieved during the first hour of the last reacquisition session. There were no differences due to Sex or Adolescent Treatment. **F)** ALPs on RF1 and ILPs across the ten sessions of extended access (6h). Self-administration was stable across the sessions (F_3.9,82.7_=1.395; p=0.243; η_p_^2^=0.06). We observed a trend for an effect of Adolescent Treatment in the average cocaine infusions during the second half of the phase (sessions 6 to 10) (F_1,21_=3.977; p=0.059; η_p_^2^=0.16). **G)** Lever pressing in the four extinction sessions as an index of seeking incubation during forced abstinence. Lever pressing behavior was found to vary across Sessions (F_2.05,43.05_=6.618; p=0.003; η_p_^2^=0.24) and, probably driven by female VEH higher lever pressing, a Sex effect (male<female) was also detected (F_1,22_=11.607; p=0.003; η_p_^2^=0.36). The visual inspection of the graph shows that after 30 days of forced abstinence THC females seem to show a lower seeking behaviour than VEH females, but the lack of a significant interaction prevents us from doing any further analysis.

## DISCUSSION

In this work, we show that chronic exposure to THC, equivalent to a mild consumption in humans, induces a broad spectrum of alterations in brain structure and function in parallel with modifications in several psychological processes related with reward and in the behavioural and cellular responses to cocaine that, nevertheless, do not fully support the Gateway Hypothesis.

### 1. Adolescent THC exposure and the psychological processes involved in reward-related behaviour and impulsivity

The first process that we examined was pavlovian conditioned approach or sign-tracking (i.e. the tendency to approach reward-cues)/goal-tracking (i.e. the tendency to approach reward) behaviours. We observed that THC-treated rats displayed more goal-tracking behaviours than their control-littermates, irrespective of their sex. This increased goal-tracking in cannabinoid-exposed animals is consistent with data that indicate that adult exposure to the cannabinoid agonist CP 55,940 increases goal-tracking behaviour (48). It is also coherent with a prior report that showed that adolescent exposure to the CB1/CB2 receptor agonist WIN 55,512-2 altered the usual proportion of sign-tracking/goal-tracking in rats, creating an intermediate phenotype in cannabinoid-exposed animals (in which rats approached both the food-cup and the CS^+^), that was not evident in vehicle-treated rats (37). However, our data are the first to suggest that adolescent exposure to the main psychoactive cannabinoid of the cannabis plant affects pavlovian conditioned approach. The bias towards goal-tracking or sign-tracking is indicative of incentive-salience attribution and depends on CB1 receptor activity (48, 49). Given that adolescent cannabinoid exposure reduces the number of cannabinoid receptors in several areas of the brain, including the nucleus accumbens (50–53), this decrease in sign-tracking behaviour and the subsequent increase of goal-tracking behaviour that we have observed is not at odds with the proposed neurochemical changes reported in the literature.

Pavlovian-to-instrumental transfer (PIT, also known as pavlovian motivation) was potentiated in THC-males. This process is related to the potentiation of an instrumental response upon presentation of a conditioned stimulus previously associated with the reward that is contingent with the instrumental response (27, 54). To our knowledge, there are no previous studies specifically ascertaining the effects of cannabinoids on PIT. The enhanced PIT of THC-males could contribute to the increase in cocaine infusions received by the THC-males under PR. Indeed, PIT is a predictor of cocaine self-administration in rats such that stronger PIT predicts a more substantial motivational influence of conditioned stimuli on self-administration and potentiates the learning of drug-cue associations (55). The precise mechanisms of potentiated PIT in THC-males are not clear. We have not detected abnormalities (either structural –MRI- or functional –PET-, or after cocaine-exposure, see c-Fos data later on) in the NAcc (a crucial region modulating PIT (56)) of these rats, suggesting that it could be the central amygdala (57) the brain region responsible for these PIT alterations. This possibility should be tested in further studies.

We also examined habit formation tendency by using a sensory-specific satiation paradigm (58). The transition from goal-directed behaviour to a habit system has been proposed to play a critical role in the development of addiction (59). We found no differences in the tendency to form habit-like responses in our adult rats exposed to THC during adolescence. This is interesting in the general context of the involvement of the endocannabinoid systems in habit formation (60–62). We and others have shown that a regime of THC exposure in adult animals accelerated habit formation (62, 63) and, we also found that the THC treatment increased dendritic spine density in the distal part of the dendrites of medium spiny neurons in the posterior dorsomedial striatum (64) suggesting a potential mechanism of these alterations in the balance between goal-directed behaviour and habits. However, when the treatment occurred during adolescence, we observed no such behavioural effects. This is in accordance with the differential effects that cannabinoids exert in the adolescent brain as compared to the adult brain in terms of the desensitisation of cannabinoid receptors. Indeed adolescent male (but not female) rats showed decreased desensitisation of the CB1 receptors in the striatum after a chronic THC treatment as compared to adult rats (65).

Lastly, we have analysed motor impulsivity using an adaptation of the widely used 5-choice serial reaction time task for boxes with two levers instead of nose-pokes. By using this 2-CSRTT, we have observed that adult rats exposed to THC during adolescence were more impulsive than their vehicle-exposed littermate controls. This effect seemed to be stronger in the females while in the males it was only evident in the second of the three tests performed. Impulsivity is a complex construct. Different varieties of impulsivity exist such as waiting impulsivity (captured by delay discounting procedures and the 2-CSRTT for example) and stopping impulsivity (the inability to stop an action that has already been initiated) (66). Within waiting impulsivity, impulsive choice (as measured in delay discounting tasks) is potentiated in adult rats that have been exposed to cannabinoids during adolescence (67). However, there is no information on the effect of adolescent cannabinoid exposure in impulsive action, another form of waiting impulsivity that has been suggested to be a predisposing endophenotype for the development of addiction (31). The fact that this form of impulsivity is potentiated in THC-exposed rats suggested that they could be more susceptible to the development of addiction-like behaviour which was another important goal of the present experiments.

### 2. Grey and white matter volumetric and microstructure alterations

Our combined volumetric and DTI analysis showed that adolescent THC induced profound structural alterations in several brain nuclei. For example, in the dorsal striatum, female rats exposed to THC showed a reduction in the volume of this structure. This result is novel since human evidence to date has not reported dorsal striatal volumetric alterations in adults with adolescent cannabis consumption. Only two studies have examined these volumetric alterations in striatal nuclei after cannabis exposure in adolescents or young people. One study examined the long-term effects of cannabis use by young people (average age of participants: 21.3 years) on the structure of the striatum (including the NAcc, the caudate and putamen) and found no significant differences between heavy cannabis users and controls or predictive effects of the age of onset of cannabis use (average age of onset: 18.8 years) (68). Using a more subtle analysis, another study found evidence for an outward deflection in the shape of the right NAcc that was predicted by the age of onset (69), but no effects were reported for the dorsal striatum. No prior data are available regarding DTI measurements in the dorsal striatum after adolescent cannabinoid exposure. The globus pallidus was also smaller in THC-treated animals (especially in the right hemisphere). This structure is an output region of the dorsal striatum and no prior studies have documented volumetric alterations after cannabis exposure. Only one study showed that in young people with cannabis use disorder (with an age of onset of 16.7 years) as compared to controls, there were alterations in shape in the *globus pallidus*, but with no volumetric alterations (70). A decreased FA value was also observed in this nucleus.

In the septal nuclei, we detected DTI alterations in the FA (decreased in THC animals) and MD (decreased in THC-males) pointing to fibre integrity alterations while in the thalamus we observed increased FA values in THC-males. We would like to highlight that the interpretation of DTI measurements is complex, as these values relate to water diffusion at the microscopic level. This is affected by several tissue parameters, such as the coherence, orientation and integrity of the fibers, as well as myelination or cell densities (71). Consequently, processes such as neuronal or axonal loss, inflammation or gliosis may have concurrently affect DTI measurements. A crucial limitation of the DTI technique is its assumption of Gaussian distribution of water diffusion. As regards this, DTI measurements work adequately in regions that contain tissue architectures that are oriented homogeneously (such as white matter), but not in regions with crossing fibre orientations (72). Thus, the pattern of DTI results reported here for the grey matter should be interpreted as a marker of THC exposure, rather than indicative of a particular histopathological phenomenon.

We have also found evidence for decreased ventricular volume (most likely due to a decrease in the volume of the lateral ventricles, specifically) in THC-exposed animals. Moreover, there was a lower aqueduct volume in male (but not female) rats exposed to THC. The mechanisms involved in this ventricular volume reductions are not clear, but acute THC injections have been shown to reduce the production of cerebrospinal fluid (CSF) (73), and THC is actively accumulated by the choroid plexus epitelium (74); perhaps the chronic THC treatment administered might have affected the basic mechanisms of CSF production and circulation thus reducing the ventricular volume.

Concerning the white matter analyses, we have found decreased FA values in the rostral anterior commissure and the rostral corpus callosum in THC-exposed rats and the hippocampal commissure in THC-exposed female rats. Decreased FA values in tracts are more straightforward to interpret than in grey matter (see above) and have traditionally been regarded as to reflect a loss of white matter integrity (75). Converging data in humans show that adolescent cannabis consumption is also linked to decreased FA in the corpus callosum (76, 77). The hippocampal commissure, an integral part of the fornix, is also compromised in cannabis users. (78). However, this is the first time that microstructural alterations in the anterior commissure are reported. In general, the alterations in the microstructure of these fibres could point to impairments in interhemispheric information processing and, especially in the case of the anterior commissure, in recognition memory (75), which is affected after adolescent cannabinoid exposure (7, 8).

### 3. Metabolic and functional alterations (^1^H-MRS and PET studies)

We detected a reduction in the levels of choline compounds in the cortex of THC-exposed rats. To the best of our knowledge, there are no previous data on GPCh+PCh alterations in the cortex after cannabis consumption or cannabinoid exposure. However, there is one report that observed decreased choline levels in cannabis users in the basal ganglia (79). Choline-containing compounds play an essential role in the synthesis and degradation of cellular membranes and are considered sensitive to changes in membrane turnover (80). The functional implications of the decrease in choline compounds are not entirely clear, but they may be indicative of an abnormal lengthening of the normal myelination dynamics that occur through postnatal and adolescent development or a neurodevelopmental delay as a result of reduced glial cell density (81).

Our preliminary developmental PET study showed maturational effects in all the ROIs analysed, and interesting sex differences in the hippocampus and the caudate nucleus.Also, our SPM analysis of the adult brain suggested a hypometabolism in a cluster of voxels comprising the inferior colliculus and the cerebellum, as well as the motor and sensory cortices of THC-exposed females. To the best of our knowledge, there are no long-term PET studies with [^18^F]-FDG in humans that have ascertained the functional effects of adolescent cannabis use. However, two previous reports by our group suggested that adolescent exposure to the synthetic cannabinoid CP 55,940 modifies brain metabolism in the frontal and amygdalo-entorhinal cortices in females (82). Moreover, the brain responses to a cocaine injection were also different in animals exposed to CP 55,940 during adolescence (83). Although these results should be replicated, they represent the first PET evidence indicative of a long-lasting alteration in the cerebellum. However, prior studies had already documented substantial cellular alterations in the cerebellum after exposure to THC. The mechanisms of these alterations are beginning to be unveiled and likely involve microglial activation. In mice, for example, subchronic administration of THC activated cerebellar microglia and increased the expression of neuroinflammatory markers, including IL-1β. Moreover, this neuroinflammatory phenotype correlated with deficits in cerebellar conditioned learning and fine motor coordination (84). The sensorimotor cortex was also affected, specifically in the females as well as the hippocampus and, to a lesser extent, the inferior colliculus. These functional alterations could also be involved in the long-term cognitive effects of adolescent cannabis use (7, 85) either alone, as separate structures, or in combination, as networks. Indeed, the cerebellum receives indirect connections from the sensory cortices to sensory information used for spatial representation (86), and there is a hippocampo-cerebellar centred network for the learning and execution of sequencebased navigation (87). There are multiple reports of altered spatial learning and memory deficits in rodents after adolescent exposure to cannabinoids (7), and the metabolic alterations preliminarily reported here could provide a mechanistic explanation.

### 4. A transcriptomic analysis of gene expression patterns in the nucleus accumbens

Following our extensive behavioural and neural characterisation of the longterm consequences of adolescent THC exposure, we focused on the nucleus accumbens, a critical region mediating the acute reinforcing effects of drugs of abuse. We performed a transcriptomic characterisation of the gene expression networks altered by the cannabinoid treatment. There were approximately the same number of DEGs for male and females but with minimal overlap (96 DEGs in the males and 87 in the males, with only 9 genes being shared between both sexes).

In the males, the gene ontology with a higher degree of representation was ‘behaviour’ with sub-categories like ‘adult locomotor behaviour’ or ‘adult walking behaviour’. In addition to their relation with behavioural processes, this ontology comprised genes related to neurotransmission and the electrophysiology of the cell, and performed important functions such as glutamate transport (*Slc1a2*) and potassium-chloride cotransport (*Slc12a5*). There were also genes coding for subunits of potassium inwardly rectifying channels (*Kcnj10*), calcium voltage-gated channels (*Cacna1e*) and the calcium-binding protein calbindin (*Calb1*), to mention some of the most representative examples. These alterations are coherent with another ontology that was modulated, ‘acidic aminoacid transport’ and also ‘neurotransmitter transport’. The alteration in the expression of these genes could alter the general physiology of nucleus accumbens cells with important consequences for drug-induced reward and motivation and also learning and memory processes, as suggested by the ‘learning and memory’ ontology that was also affected by the adolescent THC treatment.

There were also two genes belonging the top 5 DEGs with higher fold change (either up-regulated or down-regulated) that were present in the ontologies modified in the males. These two genes were *Satb2* (SATB homeobox 2, up-regulated) and *Notch3* (notch receptor 3, down-regulated). An elegant prior study has suggested that CB1 receptors are coupled to the regulation of the Ctip2–Satb2 transcriptional regulatory code (88), and in so doing, they regulate corticospinal motor neuron differentiation. The alteration of the *Satb2* gene in the nucleus accumbens of our animals could also have developmental consequences in the morphology or function of accumbal neurons as suggested by the ontologies affected by THC treatment. Satb2 in the paraventricular thalamus is also sensitive to cocaine rewarding actions (89), so it could be speculated that this up-regulated gene in accumbal cells could increase the rewarding actions of cocaine, as observed in the males after adolescent THC exposure (see below). The notch signalling pathway is also involved in brain development (90). More specifically, the notch 3 receptor (down-regulated in THC-males in these experiments) promotes neuronal differentiation, having a role opposite to Notch1/2 (91). Interestingly, the *Notch3* has also been shown to be down-regulated in striatal territories in spontaneously hypertensive rats treatred with methylphenidate during adolescence (and that further self-administered methylphenidate as adults) (92), suggesting that this gene is responsive to several pharmacological challenges during adolescence (not just cannabinoids) and also that its down-regulation may predispose to psychostimulant consumption.

In the females, the main ontology affected by THC exposure during adolescence was ‘cilium or flagellum dependent cell motility’ with three of the top-five down-regulated genes (*Aurkb*, *Ak7* and *Cfap44*) present in it. Cilia are essential structures for cell motility and migration and are also involved in several (neuro)developmental diseases (93). In addition, they are also involved in hippocampal function and cortical development (93). A significant modification of the genes belonging to this ontology by adolescent THC may point to an altered maturation of the nucleus accumbens. However, the cilliar involvement in adolescent brain development is currently unknown and clearly deserves further research. In addition to this ontology, we also obtained a hit for the ‘dicarboxylic acid transport’ category, with the *Gal* and *Folr1* genes (one of the top 5 up-regulated and down-regulated genes, respectively) involved. In addition to those two genes, other genes in the ontology were related to glutamate, GABA and hormonal signalling. The last category that we would like to highlight is ‘nicotine addiction’, again with alterations in glutamate and GABA-related genes and also adenylate cyclase 8 and nerve growth factor receptor. The fact that adolescent THC affects this category in the females is consistent with the report that adolescent cannabinoid exposure alters the rewarding properties of nicotine in adult female mice (92, but see 93). In addition, this could also point to the fact that the rewarding properties of other drugs such as cocaine could also be affected, as we shall examine in the next section of this Discussion.

### 5. Adolescent THC exposure effects on the acute actions of cocaine in the adult brain and on cocaine addiction-like behaviour

The cellular activation (as indicated by c-Fos levels) induced by cocaine was potentiated in the motor cortex by the THC treatment during adolescence (both in males and females). Although the motor cortex has not been traditionally considered as to belong to the circuit mediating the rewarding actions of drugs, some evidence points to its involvement in cocaine-induced reward. For example, the establishment of cocaine place preference is associated with increased c-Fos levels in the motor cortex, among other regions (96). Moreover, in cocaine abusers, there is hyperexcitability of the motor cortex (as revealed by lower reactivity thresholds of this area in response to magnetic stimulation) (97). In addition to the motor cortex, we also observed in the dorsomedial hypothalamic nucleus a trend for higher cocaine-induced c-Fos accumulation (as compared to the corresponding saline group) in THC females and also higher cocaine-associated c-Fos levels in the THC-males as compared to the cocaine group of VH-males. This increased reactivity of the dorsomedial hypothalamic neurons after cocaine could be related to several actions of the drug, from its interoceptive-cardiovascular effects (98) to the anorexigenic actions (99). These and other studies (40) suggest that the initial actions of cocaine are indeed modified by adolescent exposure to cannabinoids, which could also imply an alteration of cocaine addiction-like behaviour.

Several pieces of epidemiological and social research have suggested that cannabis may be one of the first steps of a series of progressions into the consumption of other drugs later (Gateway Hypothesis) (17, 100, 101). This proposed series of progressions have been criticised in studies coming from the human literature (9, 11, 12, 14, 100, 102, 103). Several attempts have been made using animal models to study the potential causal relationship between a prior cannabinoid exposure and the sensitivity to drugs of abuse later in life (see (7) for a review). With regard to cocaine, we have previously shown that cocaine self-administration acquisition was potentiated in female (but not male) rats after adolescence exposure to a cannabinoid agonist (23). Other authors have found similar effects with THC exposure during adolescence (22) (even in zebra finches using a conditioned place preference approach (104)) but not with WIN 55,512-2, a synthetic cannabinoid agonist (24). However, all these previous experiments (including our own) do not provide a comprehensive picture of addiction-like behaviour, so we decided to examine several cardinal features of addictive behaviour in our THC-exposed rats. In accordance with the previous studies (24), we found no differences in the acquisition of cocaine self-administration between THC- or vehicle exposed rats. We have used an intermediate dose of cocaine (0.5 mg/kg) which may explain the divergence between our results and those of Friedman and colleagues (22) who found potentiated cocaine self-administration with lower doses (0.1 mg/kg). In spite of this, THC-exposed male (but not female) rats earned a higher number of cocaine infusions under progressive ratio, suggesting a higher motivation for the drug. This high motivation is considered one of the hallmarks of addiction-like behaviour (31). Contrarily to this pattern, we found that after the progressive ratio phase, when rats were rebaselined in FR1 conditions, THC-exposed female (but not male) rats showed a more potent rebound responding than their vehicle-exposed littermates. The compulsive taking of the drug is another cardinal feature of addiction-like behaviour and has been typically evaluated using punished cocaine self-administration procedures (25, 31). In our experiments, all the rats diminished their cocaine intake in a similar proportion, suggesting that THC during adolescence does not potentiate compulsive cocaine intake in rats. Lastly, there was no increased self-administration under exceded access conditions or potentiated incubation of cocaine seeking as a result of THC exposure during adolescence, suggesting that adolescent THC does not increase the risk for loss of control over consumption after extended cocaine use or relapse (at least not the one precipitated by conditioned cues).

### 6. Concluding remarks and future research

In this work, we have carried out a wide-scope and multiparametric evaluation of the long-term consequences of cannabinoid exposure in the developing adolescent brain, with particular emphasis on reward processing and the potential of cannabis to act as a gateway drug leading to cocaine addiction later in life. Our results suggest that there is a causal relationship between the exposure to THC during adolescence and several alterations in brain structure and function and with a broad set of psychological alterations affecting reward processing, impulsivity and the cellular responses to cocaine, which may have resulted in higher predisposition to cocaine addiction-like behaviour at adulthood. However, when we tested this possibility, we found only partial support for this hypothesis (i.e. increased motivation for cocaine in the males and higher rebound consumption after high demand in the females, but no differences in the acquisition of cocaine self-administration, compulsive seeking, extended-access consumption or the incubation of cue-induced seeking). There are some suggestions for future research that we would like to suggest. For example, we have used FDG as a general proxy for brain function, but other ligands could also be employed to measure the number of relevant receptors, such D2/D3 dopamine receptors (with [18F]Fallypride), or the dopamine transporter (with ([^18^F]FE-PE-2I. Additional experiments aimed to increase the sample of the PET imaging data could also strengthen the conclusions drawn from this experiment. In addition, other behaviours related to cocaine addiction, such as the preference for cocaine over a natural reward or the use of second-order schedules of reinforcement could provide additional insight on the effects of adolescent THC exposure on cocaine addiction.

There are no easy answers to complex questions, but our data indicate that even a mild THC exposure during adolescence has significant consequences for the normal development of the brain and behaviour that depend on the sex of the individual. We wish to suggest that the results here presented be carefully pondered and integrated into the long-standing debate about cannabis legalisation, with an especial focus on the adolescent population.

## Acknowledgements

We would like to thank Alberto Marcos, Rosa Ferrado Luis Carrillo, Luis Troca and Gonzalo Moreno for excellent technical assistance. This manuscript was prepared during the COVID-19 pandemic and is dedicated to those who helped our society. We honour them all.

Funding provided by the Spanish Ministry of Economy and Competitiveness (PSI2016-80541-P to EA and AH-M); Spanish Ministry of Health, Social Services and Equality (Network of Addictive Disorders – Project n°: RTA-RD16/0017/0022 of the Institute of Health Carlos III to EA and Plan Nacional Sobre Drogas, Project n°: 2016I073 to EA and 2017I042 to AH-M); The General Directorate of Research of the Community of Madrid (Project n°: S-2011/BMD-2308; Program of I + D + I Activities CANNAB-CM to EA); The UNED (Plan for the Promotion of Research to EA); the European Union (Project n°: JUST/2017/AG-DRUG-806996-JUSTSO) and the BBVA Foundation (2017 Leonardo Grant for Researchers and Cultural Creators to AH-M). These agencies funded the study but had no further role in the study design; the collection, analysis and interpretation of data; the writing of the report; or the decision to submit the paper for publication. JO received funding from Instituto de Salud Carlos III; D-RM received a predoctoral fellowship granted by UNED and MU received a predoctoral fellowship awarded by the Ministry of Science and Innovation (BES-2011-043814). MLS was supported by the Ministerio de Ciencia, Innovación y Universidades, Instituto de Salud Carlos III (projects PI14/00860 and PI17/01766, and grant CPII14/00005), co-financed by European Regional Development Fund (ERDF), “A way of making Europe”, CIBERSAM, Delegación del Gobierno para el Plan Nacional sobre Drogas (2017/085), Fundación Mapfre and Fundación Alicia Koplowitz. MCV was supported by Fundación Tatiana Pérez de Guzmán el Bueno. The CNIC was supported by the Spanish Ministerio de Ciencia, Innovación y Universidades (MCIU) and the Pro-CNIC

## REFERENCES

1. Blakemore S-J. Imaging brain development: the adolescent brain. [Internet]. Neuroimage 2012;61(2):397–406.

2. Spear LP. The adolescent brain and age-related behavioral manifestations [Internet]. Neurosci Biobehav Rev 2000;24(4):417–463.

3. Paus T, Keshavan M, Giedd JN. Why do many psychiatric disorders emerge during adolescence?. Nat. Rev. Neurosci. 2008;9(12):947–957.

4. EMCDA. European Drug Report [Internet]. 2019:

5. Hurd YL. Cannabis and the developing brain challenge risk perception [Internet]. J. Clin. Invest. 2020;130(8). doi:10.1172/jci139051

6. Ferland JMN, Hurd YL. Deconstructing the neurobiology of cannabis use disorder [Internet]. Nat. Neurosci. 2020;23(5):600–610.

7. Higuera-Matas A, Ucha M, Ambrosio E. Long-term consequences of perinatal and adolescent cannabinoid exposure on neural and psychological processes [Internet]. Neurosci. Biobehav. Rev. 2015;55:119–146.

8. Stringfield SJ, Torregrossa MM. Disentangling the lasting effects of adolescent cannabinoid exposure [Internet]. Prog. Neuro-Psychopharmacology Biol. Psychiatry 2021;104:110067.

9. Mayet A, Legleye S, Beck F, Falissard B, Chau N. The gateway hypothesis, common liability to addictions or the route of administration model a modelling process linking the three theories. Eur. Addict. Res. 2016;22(2):107–117.

10. Fergusson DM, Boden JM, Horwood LJ. Cannabis use and other illicit drug use: testing the cannabis gateway hypothesis [Internet]. Addiction 2006;101(4):556–569.

11. Vanyukov MM et al. Common liability to addiction and “gateway hypothesis”: Theoretical, empirical and evolutionary perspective. Drug Alcohol Depend. 2012;123(SUPPL. 1). doi:10.1016/j.drugalcdep.2011.12.018

12. Kleinig J. Ready for Retirement: The Gateway Drug Hypothesis. Subst. Use Misuse 2015;50(8–9):971–975.

13. Nkansah-Amankra S, Minelli M. “Gateway hypothesis” and early drug use: Additional findings from tracking a population-based sample of adolescents to adulthood. Prev. Med. Reports 2016;4:134–141.

14. Tarter RE, Vanyukov M, Kirisci L, Reynolds M, Clark DB. Predictors of marijuana use in adolescents before and after licit drug use: Examination of the gateway hypothesis. Am. J. Psychiatry 2006;163(12):2134–2140.

15. Kandel DB. Does marijuana use cause the use of other drugs? [Internet]. JAMA 2003;289(4):482–483.

16. Lynskey MT, Agrawal A. Denise Kandel’s classic work on the gateway sequence of drug acquisition. Addiction [published online ahead of print: 2018]; doi:10.1111/add.14190

17. Kandel DB, Yamaguchi K, Chen K. Stages of progression in drug involvement from adolescence to adulthood: Further evidence for the gateway theory. J. Stud. Alcohol 1992;53(5):447–457.

18. Biscaia M et al. Sex-dependent effects of periadolescent exposure to the cannabinoid agonist CP-55,940 on morphine self-administration behaviour and the endogenous opioid system. Neuropharmacology 2008;54(5). doi:10.1016/j.neuropharm.2008.01.006

19. Ellgren M, Spano SM, Hurd YL. Adolescent Cannabis Exposure Alters Opiate Intake and Opioid Limbic Neuronal Populations in Adult Rats [Internet]. Neuropsychopharmacology 2007;32(3):607–615.

20. Tomasiewicz HC et al. Proenkephalin mediates the enduring effects of adolescent cannabis exposure associated with adult opiate vulnerability. Biol. Psychiatry 2012;72(10):803–810.

21. Nguyen JD, Creehan KM, Kerr TM, Taffe MA. Lasting effects of repeated Δ9-tetrahydrocannabinol vapour inhalation during adolescence in male and female rats. Br. J. Pharmacol. 2020;177(1):188–203.

22. Friedman AL, Meurice C, Jutkiewicz EM. Effects of adolescent Δ9-tetrahydrocannabinol exposure on the behavioral effects of cocaine in adult Sprague-Dawley rats. Exp. Clin. Psychopharmacol. 2019;27(4):326–337.

23. Higuera-Matas A et al. Augmented acquisition of cocaine self-administration and altered brain glucose metabolism in adult female but not male rats exposed to a cannabinoid agonist during adolescence. [Internet]. Neuropsychopharmacology 2008;33(4):806–813.

24. Kononoff J et al. Adolescent cannabinoid exposure induces irritability-like behavior and cocaine cross-sensitisation without affecting the escalation of cocaine self-administration in adulthood [Internet]. Sci. Rep. 2018;8(1):13893.

25. Deroche-Gamonet V, Belin D, Piazza P V. Evidence for addiction-like behavior in the rat [Internet]. Science (80-.). 2004;305(5686):1014–1017.

26. Everitt BJ, Giuliano C, Belin D. Addictive behaviour in experimental animals: Prospects for translation. Philos. Trans. R. Soc. B Biol. Sci. 2018; doi:10.1098/rstb.2017.0027

27. Cartoni E, Balleine B, Baldassarre G. Appetitive Pavlovian-instrumental Transfer: A review. Neurosci. Biobehav. Rev. 2016;71:829–848.

28. Fitzpatrick CJ, Morrow JD. Pavlovian Conditioned Approach Training in Rats. [Internet]. J. Vis. Exp. 2016;2016(108):e53580.

29. Everitt BJ, Robbins TW. From the ventral to the dorsal striatum: Devolving views of their roles in drug addiction [Internet]. Neurosci. Biobehav. Rev. 2013;37(9):1946–1954.

30. Belin D, Belin-Rauscent A, Murray JE, Everitt BJ. Addiction: failure of control over maladaptive incentive habits. [Internet]. Curr. Opin. Neurobiol. 2013;23(4):564–72.

31. Belin D, Mar AC, Dalley JW, Robbins TW, Everitt BJ. High impulsivity predicts the switch to compulsive cocaine-taking. Science (80-.). 2008;320(5881):1352–1355.

32. Jupp B, Dalley JW. Behavioral endophenotypes of drug addiction: Etiological insights from neuroimaging studies [Internet]. Neuropharmacology 2014;76(PART B):487–497.

33. Tomasiewicz HC et al. Proenkephalin mediates the enduring effects of adolescent cannabis exposure associated with adult opiate vulnerability. Biol. Psychiatry 2012;72(10):803–810.

34. Biscaia M et al. Sex-dependent effects of periadolescent exposure to the cannabinoid agonist CP-55,940 on morphine self-administration behaviour and the endogenous opioid system. Neuropharmacology 2008;54(5):863–873.

35. Ellgren M et al. Dynamic changes of the endogenous cannabinoid and opioid mesocorticolimbic systems during adolescence: THC effects. Eur. Neuropsychopharmacol. 2008;18(11):826–834.

36. Higuera-Matas A et al. Sex-specific disturbances of the glutamate/GABA balance in the hippocampus of adult rats subjected to adolescent cannabinoid exposure [Internet]. Neuropharmacology 2012;62(5–6):1975–1984.

37. Schoch H et al. Adolescent cannabinoid exposure effects on natural reward seeking and learning in rats. Psychopharmacology (Berl). 2017;1–14.

38. Higuera-Matas A et al. Periadolescent exposure to cannabinoids alters the striatal and hippocampal dopaminergic system in the adult rat brain. [Internet]. Eur. Neuropsychopharmacol. 2010;20(12):895–906.

39. Miller ML et al. Adolescent exposure to Δ 9 - tetrahydrocannabinol alters the transcriptional trajectory and dendritic architecture of prefrontal pyramidal neurons [Internet]. Mol. Psychiatry doi:10.1038/s41380-018-0243-x

40. Scherma M et al. Cannabinoid exposure in rat adolescence reprograms the initial behavioral, molecular, and epigenetic response to cocaine. Proc. Natl. Acad. Sci. U. S. A. 2020;117(18):9991–10002.

41. Chye Y, Christensen E, Yücel M. Cannabis Use in Adolescence: A Review of Neuroimaging Findings [Internet]. J. Dual Diagn. 2019;0(0):1–23.

42. Ganzer F, Bröning S, Kraft S, Sack PM, Thomasius R. Weighing the Evidence: A Systematic Review on Long-Term Neurocognitive Effects of Cannabis Use in Abstinent Adolescents and Adults. Neuropsychol. Rev. 2016;26(2):186–222.

43. Higuera-Matas A et al. Chronic cannabinoid administration to periadolescent rats modulates the metabolic response to acute cocaine in the adult brain. Mol. Imaging Biol. 2011;13(3):411–5.

44. Bari A, Dalley JW, Robbins TW. The application of the 5-choice serial reaction time task for the assessment of visual attentional processes and impulse control in rats. [Internet]. Nat. Protoc. 2008;3(5):759–67.

45. Kallio MA et al. Chipster: User-friendly analysis software for microarray and other high-throughput data. BMC Genomics 2011;12(507). doi:10.1186/1471-2164-12-507

46. Zhou Y et al. Metascape provides a biologist-oriented resource for the analysis of systems-level datasets. Nat. Commun. 2019;10(1):1523.

47. Bodineau L et al. The c-FOS Protein Immunohistological Detection: A Useful Tool As a Marker of Central Pathways Involved in Specific Physiological Responses In Vivo and Ex Vivo. J. Vis. Exp. 2016;(110):1–9.

48. Gheidi A et al. Effects of the cannabinoid receptor agonist CP-55,940 on incentive salience attribution [Internet]. Psychopharmacology (Berl). [published online ahead of print: 2020]; doi:10.1007/s00213-020-05571-3

49. Bacharach SZ et al. Cannabinoid receptor-1 signaling contributions to sign-tracking and conditioned reinforcement in rats. Psychopharmacology (Berl). 2018;235(10):3031–3043.

50. Higuera-Matas A et al. Sex-specific disturbances of the glutamate/GABA balance in the hippocampus of adult rats subjected to adolescent cannabinoid exposure. Neuropharmacology 2012;62(5–6). doi:10.1016/j.neuropharm.2011.12.028

51. Zamberletti E et al. Gender-dependent behavioral and biochemical effects of adolescent delta-9-tetrahydrocannabinol in adult maternally deprived rats. Neuroscience 2012;204:245–257.

52. Rubino T et al. Chronic Delta(9)-Tetrahydrocannabinol During Adolescence Provokes Sex-Dependent Changes in the Emotional Profile in Adult Rats: Behavioral and Biochemical Correlates. Neuropsychopharmacology 2008;

53. Kruse LC, Cao JK, Viray K, Stella N, Clark JJ. Voluntary oral consumption of Δ 9 - tetrahydrocannabinol by adolescent rats impairs reward-predictive cue behaviors in adulthood[published online ahead of print: 2019];(April). doi:10.1038/s41386-019-0387-7

54. Corbit LH, Balleine BW. Learning and motivational processes contributing to pavlovian–instrumental transfer and their neural bases: Dopamine and beyond. Curr. Top. Behav. Neurosci. 2016;27:259–289.

55. Takahashi TT, Vengeliene V, Enkel T, Reithofer S, Spanagel R. Pavlovian to instrumental transfer responses do not correlate with addiction-like behavior in rats. Front. Behav. Neurosci. 2019;13. doi:10.3389/fnbeh.2019.00129

56. Corbit LH, Balleine BW. The general and outcome-specific forms of Pavlovian-instrumental transfer are differentially mediated by the nucleus accumbens core and shell. [Internet]. J. Neurosci. 2011;31(33):11786–94.

57. Corbit LH, Balleine BW. Double dissociation of basolateral and central amygdala lesions on the general and outcome-specific forms of pavlovian-instrumental transfer. [Internet]. J. Neurosci. 2005;25(4):962–70.

58. Hogarth L, Balleine BW, Corbit LH, Killcross S. Associative learning mechanisms underpinning the transition from recreational drug use to addiction. Ann. N. Y. Acad. Sci. 2013;1282(1):12–24.

59. Everitt BJ, Robbins TW. Drug Addiction: Updating Actions to Habits to Compulsions Ten Years On. [Internet]. Annu. Rev. Psychol. [published online ahead of print: August 7, 2015]; doi:10.1146/annurev-psych-122414-033457

60. Hilário MRF, Clouse E, Yin HH, Costa RM. Endocannabinoid signaling is critical for habit formation. [Internet]. Front. Integr. Neurosci. 2007;1(November):6.

61. Gremel CM et al. Endocannabinoid Modulation of Orbitostriatal Circuits Gates Habit Formation [Internet]. Neuron 2016;1–13.

62. Nazzaro C et al. SK channel modulation rescues striatal plasticity and control over habit in cannabinoid tolerance. [Internet]. Nat. Neurosci. 2012;15(2):284–93.

63. Fernández-Cabrera M, Ucha M, Ambrosio E, Miguéns M, Higuera-Matas A. A Chronic Treatment with Δ-9 Tetrahydrocannabinol Accelerates Habit Formation in Mice. In: XVI Reunión de la Sociedad Española de Investigación sobre Cannabinoides. Cuenca: 2014:

64. Fernández-Cabrera MR et al. Selective effects of Δ9-tetrahydrocannabinol on medium spiny neurons in the striatum. PLoS One 2018;13(7). doi:10.1371/journal.pone.0200950

65. Burston JJ, Wiley JL, Craig AA, Selley DE, Sim-Selley LJ. Regional enhancement of cannabinoid CB 1 receptor desensitisation in female adolescent rats following repeated Δ 9-tetrahydrocannabinol exposure. Br. J. Pharmacol. 2010;161(1):103–112.

66. Dalley JW, Robbins TW. Fractionating impulsivity: Neuropsychiatric implications. Nat. Rev. Neurosci. 2017;18(3):158–171.

67. Johnson KR, Boomhower SR, Newland MC. Behavioral effects of chronic WIN 55,212-2 administration during adolescence and adulthood in mice. Exp. Clin. Psychopharmacol. 2019;27(4):348–358.

68. Cousijn J et al. Grey matter alterations associated with cannabis use: Results of a VBM study in heavy cannabis users and healthy controls [Internet]. Neuroimage 2012;59(4):3845–3851.

69. Orr JM, Paschall CJ, Banich MT. Recreational marijuana use impacts white matter integrity and subcortical (but not cortical) morphometry [Internet]. NeuroImage Clin. 2016;12:47–56.

70. Smith MJ et al. Cannabis-related working memory deficits and associated subcortical morphological differences in healthy individuals and schizophrenia subjects [Internet]. Schizophr. Bull. 2014;40(2):287–299.

71. Antonsen BT et al. Altered diffusion tensor imaging measurements in aged transgenic Huntington disease rats [Internet]. Brain Struct. Funct. 2013;218(3):767–778.

72. Beaulieu C. The basis of anisotropic water diffusion in the nervous system - A technical review [Internet]. NMR Biomed. 2002;15(7–8):435–455.

73. Mancall AC, DiGregorio GJ, Brill CB, Ruch E. The Effect of Δ-9-Tetrahydrocannabinol on Rat Cerebrospinal Fluid [Internet]. Arch. Neurol. 1985;42(11):1069–1071.

74. Agnew WF, Rumbaugh CL, Cheng JT. The uptake of Δ9-tetrahydrocannabinol in choroid plexus and brain cortex in vitro and in vivo [Internet]. Brain Res. 1976;109(2):355–366.

75. DeLisi LE. The effect of cannabis on the brain: Can it cause brain anomalies that lead to increased risk for schizophrenia? [Internet]. Curr. Opin. Psychiatry 2008;21(2):140–150.

76. Bava S, Jacobus J, Thayer RE, Tapert SF. Longitudinal Changes in White Matter Integrity Among Adolescent Substance Users [Internet]. Alcohol. Clin. Exp. Res. 2013;37:E181–E189.

77. Becker MP, Collins PF, Lim KO, Muetzel RL, Luciana M. Longitudinal changes in white matter microstructure after heavy cannabis use [Internet]. Dev. Cogn. Neurosci. 2015;16:23–35.

78. Zalesky A et al. Effect of long-term cannabis use on axonal fibre connectivity [Internet]. Brain 2012;135(7):2245–2255.

79. Chang L, Cloak C, Yakupov R, Ernst T. Combined and independent effects of chronic marijuana use and HIV on brain metabolites. [Internet]. J. Neuroimmune Pharmacol. 2006;1(1):65–76.

80. Gujar SK, Maheshwari S, Björkman-Burtscher I, Sundgren PC. Magnetic resonance spectroscopy. [Internet]. J. Neuroophthalmol. 2005;25(3):217–26.

81. Ford TC, Crewther DP. A comprehensive review of the 1H-MRS metabolite spectrum in autism spectrum disorder [Internet]. Front. Mol. Neurosci. 2016;9(MAR):14.

82. Higuera-Matas A et al. Augmented acquisition of cocaine self-administration and altered brain glucose metabolism in adult female but not male rats exposed to a cannabinoid agonist during adolescence. Neuropsychopharmacology 2008;33(4). doi:10.1038/sj.npp.1301467

83. Higuera-Matas A et al. Chronic cannabinoid administration to periadolescent rats modulates the metabolic response to acute cocaine in the adult brain. Mol. Imaging Biol. 2011;13(3). doi:10.1007/s11307-010-0388-8

84. Cutando L et al. Microglial activation underlies cerebellar deficits produced by repeated cannabis exposure. [Internet]. J. Clin. Invest. 2013;123(7):2816–31.

85. Meier MH et al. Persistent cannabis users show neuropsychological decline from childhood to midlife. Proc. Natl. Acad. Sci. U. S. A. 2012;109(40):E2657–64.

86. Rondi-Reig L, Paradis AL, Lefort JM, Babayan BM, Tobin C. How the cerebellum may monitor sensory information for spatial representation. Front. Syst. Neurosci. 2014;8(November). doi:10.3389/fnsys.2014.00205

87. Babayan BM et al. A hippocampo-cerebellar centred network for the learning and execution of sequence-based navigation. Sci. Rep. 2017;7(1). doi:10.1038/s41598-017-18004-7

88. Diaz-Alonso J et al. The CB(1) cannabinoid receptor drives corticospinal motor neuron differentiation through the Ctip2/Satb2 transcriptional regulation axis [Internet]. J. Neurosci. 2012;32(47):16651–16665.

89. Salti A et al. Cocaine paired environment increases SATB2 levels in the rat paraventricular thalamus [Internet]. Front. Behav. Neurosci. 2018;12. doi:10.3389/fnbeh.2018.00224

90. Androutsellis-Theotokis A et al. Notch signalling regulates stem cell numbers in vitro and in vivo [Internet]. Nature 2006;442(7104):823–826.

91. Rusanescu G, Mao J. Notch3 is necessary for neuronal differentiation and maturation in the adult spinal cord [Internet]. J. Cell. Mol. Med. 2014;18(10):2103–2116.

92. dela Peña I et al. Prefrontal cortical and striatal transcriptional responses to the reinforcing effect of repeated methylphenidate treatment in the spontaneously hypertensive rat, animal model of attention-deficit/hyperactivity disorder (ADHD) [Internet]. Behav. Brain Funct. 2014;10(1). doi:10.1186/1744-9081-10-17

93. Louvi A, Grove EA. Cilia in the CNS: The quiet organelle claims center stage [Internet]. Neuron 2011;69(6):1046–1060.

94. Dukes AJ, Fowler JP, Lallai V, Pushkin AN, Fowler CD. Adolescent Cannabinoid and Nicotine Exposure Differentially Alters Adult Nicotine Self-Administration in Males and Females [Internet]. Nicotine Tob. Res. 2020;22(8):1364–1373.

95. Flores Á, Maldonado R, Berrendero F. THC exposure during adolescence does not modify nicotine reinforcing effects and relapse in adult male mice [Internet]. Psychopharmacology (Berl). 2020;237(3):801–809.

96. Soderman AR, Unterwald EM. Cocaine reward and hyperactivity in the rat: Sites of mu opioid receptor modulation [Internet]. Neuroscience 2008;154(4):1506–1516.

97. Hanlon CA et al. A comprehensive study of sensorimotor cortex excitability in chronic cocaine users: Integrating TMS and functional MRI data [Internet]. Drug Alcohol Depend. 2015;157:28–35.

98. Stotz-Potter EH, Willis LR, DiMicco JA. Muscimol acts in dorsomedial but not paraventricular hypothalamic nucleus to suppress cardiovascular effects of stress. [Internet]. J. Neurosci. 1996;16(3):1173–9.

99. Bellinger LL, Bernardis LL. The dorsomedial hypothalamic nucleus and its role in ingestive behavior and body weight regulation: lessons learned from lesioning studies. [Internet]. Physiol. Behav. 2002;76(3):431–42.

100. Fergusson DM, Boden JM, Horwood LJ. Cannabis use and other illicit drug use: Testing the cannabis gateway hypothesis. Addiction 2006;101(4):556–569.

101. Yamaguchi K, Kandel DB. Patterns of drug use from adolescence to young adulthood: II. Sequences of progression. Am. J. Public Health 1984;74(7):668–672.

102. Morral AR, McCaffrey DF, Paddock SM. Reassessing the marijuana gateway effect. Addiction 2002;97(12):1493–1504.

103. Degenhardt L et al. Evaluating the drug use “gateway” theory using cross-national data: Consistency and associations of the order of initiation of drug use among participants in the WHO World Mental Health Surveys. Drug Alcohol Depend. 2010;108(1–2):84–97.

104. Aldhafiri A, Dodu JC, Alalawi A, Emadzadeh N, Soderstrom K. Delta-9-THC exposure during zebra finch sensorimotor vocal learning increases cocaine reinforcement in adulthood. [Internet]. Pharmacol. Biochem. Behav. 2019;172764.

