## Supplementary online material for "Testing the Role of Δ^9^-Tetrahydrocannabinol During Adolescence as a Gateway Drug: Behavioural, Brain Imaging and Transcriptomic Studies"

**SUPPLEMENTARY ON-LINE MATERIALS (SOM)**

**SUPPLEMENTARY METHODS**

**ANIMALS**

Wistar albino rats from Charles-River S.A. (Saint-Germain-sur-l’Arbresle, France) were mated (one male x one female) in our laboratory two weeks after their arrival, and their male and female offspring were used. On the day of birth (postnatal day –PND-0), the litters were sex-balanced and culled to a litter size of 10 ± 2 pups per dam. The animals were weaned at PND 22 and housed in plexiglass cages (2 or 3 sibling animals of the same sex and treatment per cage). All animals were maintained at a constant temperature (20±2ºC) under a reverse 12-h/12-h dark/light cycle (lights on at 20:00 h), with free access to water and food (commercial diet for rodents SAFE. Augy, France), unless otherwise specified. Different sets of animals, obtained from six different cohorts (35 litters) were randomly assigned to each of the six experiments described below, thus minimising litter effects. Every attempt was made to minimise the pain and discomfort of the experimental animals, and all procedures were conducted in accordance with the European Union legislation on the protection of animals used for scientific purposes (2010/63/EU Directive) and approved by the Ethics Board of the National University of Distance Learning.

**Δ^9^-TETRAHYDROCANNABINOL EXPOSURE DURING ADOLESCENCE**

Δ^9^-Tetrahydrocannabinol (THC) was purchased from THCPharm (Frankfurt, Germany) as resin, dissolved in pure ethanol (Merck) and stored at -30ºC in 0.5 mL aliquots in siliconised (Sigmacote; Merck) vials with a nitrogen-saturated atmosphere, protected from the light. Vehicle aliquots were treated in the same way, but no THC was added. Each day the final solution was prepared by adding to each aliquot, kolliphor (PEG-35 castor oil; Merk) and saline (0.9% NaCl solution; Vitulia, Spain) in a 1:1:18 proportion to a final volume of 10 mL. The ethanol concentration in both the THC and vehicle solution was 5% v/v, a dose of approximately 0.0789 g/kg that does not induce significant behavioural effects (1). The adolescent chronic Δ^9_^THC treatment was administered every other day from PND 28 to PND 44 (or, the developmental design implemented, from PND 38 to PND 54 in the PET experiment). Rats received an intraperitoneal injection (2 mL/kg) of Δ^9_^THC at a dose of 3 mg/kg or vehicle (2 mL/kg). Using a body surface area conversion (2) this dose is equivalent to 0.48 mg/kg in a ‘typical’ 15-year-old male adolescent (weighing 60 kg) which results in 28.8 mg of THC per exposure. This dose can be considered mild as per actual average cannabis smoking practices (for example, an amount of 320 mg of cannabis per cigarette (3) and a 13.6% THC concentration in the cannabis used (4)).

**BEHAVIOURAL STUDIES**

**Apparatus**

All the behavioural procedures (with the exception of cocaine self-administration) were performed in Med-Associates operant boxes (29.53 L x 24.84 W x 18.67 H cm) placed inside a sound-attenuating chamber equipped with a fan, a tone generator and a clicker. Rewards were food pellets (Noyes pellets; Testdiets). The food magazine placed in one of the walls was connected to a pellet dispenser located outside the box. All conditioning boxes were equipped with two retractable levers with cue lights above them in the right and left side of the food magazine.

Cocaine self-administration was performed in Coulbourn Instruments operant boxes (30 L x 25, 4 W x 30 H cm). These boxes were equipped with two levers in one of the walls, with a LED light cue over each of them and stainless-steel grid floor connected to a shock generator (Coulbourn) for the study of compulsivity. Every box was placed inside a sound-attenuating chamber and equipped with a pump (Harvard apparatus) outside of the chamber that delivered a programmed volume of the cocaine solution to the animal.

**Pavlovian to instrumental transfer & 2-Choice Serial Reaction Time Task**

Before starting each behavioural protocol (at around PND 90), animals were food-restricted, and their weight kept between 90-95% of the original in the free-feeding state.

1. PIT:

The PIT protocol started two days later. The weight gain produced by the normal development of these animals without dietary restrictions was taken as the reference for the calculation of the food regime and the weight range in which they had to be maintained. Animals received their daily food after the experimental sessions.

The PIT protocol consisted of four consecutive phases:

(1) Pavlovian training: 10 sessions of 4 cycles composed of a) 2 minutes under a variable interval of 30 s (VI30) schedule of reinforcement where they were exposed to a reward-associated conditioned stimulus (clicker or homeobox light, counterbalanced) or CS^+^; b) 1-minute trials without conditioned stimulus (Inter Stimulus Interval or ISI), where no stimuli or programmed contingencies were administered; and c) a 2-minute trial where the animals were exposed to a non-rewarded stimulus or CS^-^ (tone or clicker, counterbalanced). This is summarised as follows:

[No Stimuli Interval → CS^+/-^ → No Stimuli Interval → CS^-/+^] * 4times

During the Pavlovian training sessions of the PIT protocol, the number of head entries (HE) into the magazine under each stimulus condition was recorded, and a CS^+^ HE ratio was calculated as:

$${CS}^{+} HE ratio =\frac{{CS}^{+}HEs}{{CS}^{+}HEs +{CS}^{-}HEs}$$

(2) Instrumental training: 1 session under fixed ratio (FR) 1, three sessions under variable-ratio VR5 and three sessions under variable-ratio VR10. Sessions began with the insertion of the *active lever*, associated with the reinforcer schedule, and *inactive lever* that had no programmed consequences. All sessions were limited to 30 pellets. During the instrumental training sessions, lever presses performed on the active lever (ALP), active lever presses during time out (TOALP), inactive lever presses (ILP) and HE were recorded. To study the discrimination between the rewarded and the unrewarded lever, an ALP ratio was calculated as:

$ALP ratio=\frac{ALPs}{ALPs + ILPs}$

(3) Extinction: Two daily sessions of 20 minutes with levers protracted but no programmed consequences. On EXTINCTION SESSIONS, HE and lever presses performed on the former active or inactive levers were recorded. The total LP (ALP+ILP) was calculated.

(4) PIT test: One single session that began with a third extinction period of 20 minutes then 4 cycles composed CS or ISI presentations (counterbalanced) lasting two minutes, as indicated below:

[No Stimuli Interval → CS^+/-^ → No Stimuli Interval → CS^-/+^] * 4times

In the PIT test, HE and lever presses on former active or inactive levers under the different stimulus conditions were registered. The Pavlovian to instrumental transfer phenomenon was calculated using the percentage of ALP on CS+:

$${\%ALPs on CS}^{+}=\frac{{CS}^{+}ALPs}{{CS}^{+}ALPs+ {CS}^{-}ALPs}*100$$

We also studied the simple Pavlovian approach in contrast to the instrumental transfer, for this purpose, we calculated the percentage of HEs on CS+:

$$\%HEs on {CS}^{+}=\frac{{CS}^{+}HE}{{CS}^{+}HE+ {CS}^{-}HE}*100$$

This protocol was designed considering the factors that modulate the expression of the PIT phenomenon (5) and based on preliminary tests conducted in our laboratory.

2. 2-CSRTT:

Ten days after the end of the PIT animals were again food-restricted and the two-choice serial reaction time task (2-CSRTT) protocol (modified from the one published by Bari and Robins(6)) began. First, the rats went through two sessions of cue-lever training, in which one of the cue lights over one of the levers (right or left) remained on and lever presses on this lever were rewarded. Twelve phases with increasing demands were implemented. In each phase, the duration of a light stimulus over a lever that signalled the availability of a pellet was progressively shortened (phase 1: 30 s – phase 12: 0.5 s). Also, the response time that the rats had to press the lever in order to obtain the reward was progressively shortened (phase 1: 30 s – phase 12: 5 s). The Inter-trial-interval (ITI) time also increased across sessions (phase 1: 2 s – phase 12: 5 s). Rats progressed to the next phase if they managed to perform at least an 80% of correct lever presses (CLP), i.e. press the lever closer to the light stimulus. Further lever presses after a CLP during the response time were registered as perseverative responses (PerR), while responses in any of the levers during ITI were counted as premature responses (PreR). PreR and lever presses in the not-signalled lever, namely incorrect lever presses (ILP), or failure to respond in a trial (Omission Response-OR) were punished with a time out (TO) of 5 s. During this TO, responses were considered as time-out responses (TOR) and caused to TO period to be reinstated. Animals quickly learned to avoid responses before cues were present in order to obtain a new pellet. Once phase 12 was reached, 6 more consecutive sessions with the same requirements (but 75% of correct responses required) were implemented to serve as a baseline (BL). Then three long-ITI (9 s) sessions with two phase 12 sessions amid them, were performed.

We calculated the percentage of PreR in each BL sessions or long-ITI test session as follows:

$$\% PreR =\frac{PreR}{(PreR)+(CLP)+(ILP)+(OR)}*100$$

We additionally calculated the percentage of PreR increase in each of these long-ITI sessions as follows:

$$\% increase =\frac{PreR on test - PreR on previous BL sessions}{PreR on test}*100$$

**Pavlovian conditioned approach (sign-tracking/goal-tracking) & habit formation**

1. PCA:

At approximately PND 90, the same food restriction described previously for the PIT protocol was applied to a different subset of animals. After reaching the desired body weight, a single session of magazine training was performed before the commencement of the PCA protocol. In this session, the feeder dispensed 25 food pellets, under a variable interval of 90 s programme, into the magazine. On the following day, the animals began the PCA protocol proper, consisting of 8 daily sessions. Each session consisted of 25 trials in which the feeder dispensed pellets into the magazine under a variable interval 60 s schedule of reinforcement. A lever on one of the sides of the magazine (right or left, counterbalanced) was extended 8 s before the reward. This lever was retracted in the moment of reward presentation. The other lever was present during the whole session and served as a measure of general locomotor activity. None of the levers had programmed contingencies. Interaction with the CS^+^ lever (CS^+^ LPs), presses in the inactive lever in between each CS^+^ (ILPs) or during the CS^+^ (CS^+^ ILPs), HE and time spent into the magazine (MAG) in between each CS^+^ presentation and during the CS+ (CS^+^ HE and CS^+^ MAG, respectively) was recorded. The main PCA index used, suggestive of the bias to goal-track or sign-track was calculated each day as the mean of three other indexes:

i) Response bias, i.e., ratio of the total number of lever presses and magazine entries for a session during the CS+ presentations:

$$Response bias =\frac{{CS}^{+}LPs- {CS}^{+}HEs}{{CS}^{+}LPs+ {CS}^{+}HEs}$$

ii) Latency score, i.e., average latency to perform a lever press or magazine entry during the 8 s of CS+ presentation:

$$Latency score =\frac{Mean HEs latency- Mean LP latency}{8}$$

iii) Probability difference, i.e., the probability of performing a HE during the CS^+^ presentations minus the probability of performing a LP during the CS^+^ presentation:

$$Probability difference =\frac{LP check}{25}-\frac{HE check}{25}$$

We also calculated another additional measure: response probability (similar to probability difference but only for the first response, being a CS^+^ LP or a CS^+^ HE), performed in each trial:

$$First response probability =\frac{1st Response LP}{25}-\frac{1st Response HE}{25}$$

Every index score ranged from 1 (absolute sign-tracking) to -1 (absolute goal-tracking), with 0 representing no preference bias. Animals with a PCA score higher than 0.5 were classified as sign-trackers, whereas animals with a PCA lower than -0.5 were categorised as goal-trackers.

2. Sensory-specific devaluation (habit formation study)

Ten days after final PCA session animals began the habit training protocol. The rats performed a brief, non-habit-forming training, and an extended training scheme. Every session was implemented with a single lever protracted inside the operant box. The brief training consisted of 5 consecutive daily sessions: one FR1 session, two variable-interval (VI) 30 s sessions and two VI 60 s sessions. After the training sessions, we subjected the animals to two counterbalanced, sensory-specific satiety-based, devaluation tests**.** In these tests, the rats were allowed to eat pellets or chow food freely for one hour before undergoing an extinction test. This test was a brief extinction session of 5 minutes with two levers present, the one presented during training and a new one. This allowed us to discriminate reinforcer seeking from locomotor activity. Both levers had no programmed consequences. On the day after the first test, the animals were retrained under a regular VI60 session and, on the next day, another devaluation session was carried out with the other type of food (pellets or regular chow). Two days after the second devaluation test, the animals were again retrained on a VI60s session and performed an omission test (where the omission of the response was rewarded, i.e. every LP reset the clock that delivered the reward every 30 s) to assess their performance during contingency degradation. For extended training, animals performed another 10 VI60 sessions and then animals underwent the same counterbalanced devaluation tests and a final omission test, as described before. In order to calculate the devaluation index, we applied the following formula:

$$Habit Formation Index=\frac{{LP}_{Devalued}}{{LP}_{Devalued}+{LP}_{NonDevalued}}$$

**Magnetic Resonance Imaging**

At approximately PND 90, rats were anaesthetised with 2% isoflurane in 1 L of oxygen in an induction chamber, and the flow of anaesthetic gas regulated continuously to maintain a breathing rate of 50 ± 20 bpm (SA Instruments, Stony Brook, NY). Body temperature was maintained at approximately 37 °C using warm water through a heat exchanger blanket placed inside the animal platform. T2-weighted (T2-W) spin-echo anatomical images were then acquired in a Bruker Pharmascan 7 Tesla system (Bruker Medical Gmbh, Ettlingen, Germany) with a rapid acquisition with relaxation enhancement (RARE) sequence in axial and coronal orientations and the following parameters: TR: 3000 ms; TE: 44 ms; RARE factor: 8; Av: 3; FOV: 3.5 cm; acquisition matrix: 256×256 corresponding to an in-plane resolution of 136×136 μm^2^; slice thickness: 1.50 mm and a number of slices of 18 for axial and 8 for coronal images. Volumetric analyses were made by manually segmenting the different regions of interest (ROIs) in each image by a blind experimenter and then calculating the area with Image J. The total brain volume and relative volume of these areas (left, right and total) were calculated for the following ROIs, using anatomical landmarks within each slice for their delimitation: Striatum (STR), Nucleus Accumbens (NAcc), Hippocampus (HIPP), Cortex (Cx), Globus pallidus (GP), Thalamus (THA), Amygdala (AMY), Septal Nuclei (SNu) and Cerebellum (Cb). In addition, the volume occupied by each ventricle and total ventricular volume, both relative to total regional brain volume, were also calculated.

Diffusion-weighted images (DTI) were acquired in the same imaging session with a spin-echo single-shot echo-planar imaging (EPI) pulse sequence using the following parameters: TR/TE 3500/40ms; averages 1; diffusion gradient duration: 3.5ms; diffusion gradient separation: 20ms; gradient directions: 7; two b values (100 and 1400s/mm^2^), slices thickness 1.5 mm without a gap. All EPI data were acquired with a single-shot EPI sequence, 96x96 matrix, and zero-filled in k space to construct a 128x128 image matrix corresponding to an in-plane resolution of 273x273µm^2^. Fractional anisotropy (FA), mean diffusivity (MD), trace, the eigenvalues and eigenvector maps were calculated with a native software application written in Matlab (R2007a). The values of these indices were extracted using the Image J software from maps in selected Regions of Interest (ROIs), drawn manually in each slide, and using as reference the correspondent T2-W anatomical image and the Paxinos-Watson brain atlas (7). Grey matter values of FA and MD were obtained in the Cingulate Cortex (CCG), STR, NAcb, HIPP, GP, THA, Motor cortex and SNu. White matter FA signal was obtained similarly in the following white matter tracts: anterior commissure (ac), *corpus callosum* (cc), internal capsule (ic) and the hippocampal commissure of the fornix (fo).

In addition, following the T2 and DTI imaging protocols, an *in vivo* ^1^H magnetic resonance spectroscopy study was performed in two brain regions: cortex and striatum (one region per hemisphere, see Figure 5). The spectroscopy protocol used a Point-Resolved Spatially Spectroscopy (PRESS), combined with VAPOR water suppression and employed the following parameters: TR: 3000 ms; TE: 35 ms; Av: 128; voxel volume: 27 mm^3^. First and second-order shims were automatically adjusted with FASTMP application in a large voxel (64 mm^3^). All ^1^H spectra were automatically analysed using LCModel version 6.2-OR (Stephen Provencher, Oakville, ON; Canada). Statistical analysis was performed with the concentration values of each metabolite relative to creatinine+phosphocreatinine (Cr+PCr) for those with a standard deviation under 20% (the value of this metabolite was not affected by the *Sex* or *Adolescent Treatment* factors, or their interaction). All the magnetic resonance procedures were carried out at the SIERMAC core unit of the Alberto Sols Biomedical Research Institute (Madrid, Spain).

**Positron Emission Tomography**

PET-CT studies were performed in adult rats (between PND 65 and PND 69) at the Radioisotopes for Biomedicine research group of the Center for Energy, Environmental and Technological Research (CIEMAT) in Madrid, Spain, using a small-animal PET-CT (SEDECAL, Madrid, Spain).

PET-CT studies were performed in adult rats at PND 32-33 and PND 60, using an Argus PET/CT scanner (SEDECAL, Madrid, Spain). Briefly, the rats were anaesthetised using 2–3% isoflurane in medical oxygen (1 L/min). The temperature was maintained at 37ºC using a heating pad during PET acquisition. Static PET imaging was obtained for 45 min at 30 min post intravenous administration with 176 ± 37 MBq/kg body weight of 2-deoxy-2-[^18^F]fluoro-D-glucose (2-[^18^F]FDG). PET data were reconstructed using a 2D-OSEM algorithm (16 subsets and 3 iterations) with random and scatter corrections.

PET images underwent a pre-processing protocol previously described (4). Briefly, each PET image was spatially co-registered to a common reference CT scan for each sex by an automatic method based on mutual information (8), followed by 9 points scaling in the three spatial directions. Then, PET intensity values were normalised to the mean brain intensity. Four brain ROIs (hippocampus, prefrontal cortex, caudate nucleus and cortex) were segmented. The statistical analysis consisted in a three-way mixed analysis of variance (ANOVA), followed by Bonferroni multiple comparison post-test, with the between-subjects factors representing the gender and treatment (THC or vehicle) and the within-subjects factor representing the time. Statistical analyses were performed in SSPS 14.0. A p-value of less than 0.05 was considered statistically significant.

Also, for the PND65 scans, voxel-based 2-sample t-tests (p<0.05 uncorrected) were performed with Statistical Parametric Mapping (SPM) software (http://www.fil.ion.ucl.ac.uk/spm/software/spm12/) for each sex. PET images were smoothed with a gaussian kernel of 2.5 times the voxel size of full width at a half maximum (FWHM) and masked in order to exclude extracerebral voxels from the analyses. Only clusters more extensive than 50 adjacent voxels were considered in order to minimise the effect of type I errors.

**RNASeq transcriptomic analysis of the nucleus accumbens**

At PND 90, animals were deeply anaesthetised with isoflurane and sacrificed by decapitation. Brains were extracted and, with the help of a brain matrix, 1 mm thick coronal slices were obtained between 2.28 mm and 1.08 mm, approximately, anterior from bregma. With the help of two dissecting lancet-shaped needles, the NAcb (shell division) was dissected according to the Paxinos and Watson atlas(9). All the surfaces and tools used for dissection were sterilised and treated with RNAseZap® (Ambion), and all the steps were carried out with caution to maintain RNA integrity. The tissue samples were snap-frozen with dry ice and stored at -70ºC. Samples were homogenised, and RNA extracted according to the protocol, tools and reagents provided by RNeasy Mini Kit (Qiagen). Libraries were prepared according to the instructions of the *NEBNext Ultra Directional RNA Library Prep kit for Illumina* kit (New England Biolabs), as detailed in “Chapter 1: Protocol for use with NEBNext Poly(A) mRNA Magnetic Isolation Module”. The input yield of total RNA to start the protocol was 1 µg quantified by an Agilent 2100 Bioanalyzer using an RNA 6000 nano LabChip kit. We performed the library amplification included in the cited protocol using a PCR of 14 cycles. The obtained libraries were validated and quantified by an Agilent 2100 Bioanalyzer using a DNA7500 LabChip kit, and an equimolecular pool of libraries were titrated by quantitative PCR using the “Kapa-SYBR FAST qPCR kit forLightCycler480” (Kapa BioSystems) and a reference standard for quantification. The pool of libraries was denatured prior to being seeded on a flow-cell at a density of 2.2 pM, where clusters were formed and sequenced using a “NextSeq™ 500 High Output Kit”, in a 1x75 single read sequencing run on a NextSeq500 sequencer.

We used the Chipster analysis suite (10) to perform data processing and analysis. Briefly, data quality analysis of the obtained raw data (singleEnd, stranded) was performed on FASTQC and PRINSEQ; no low-quality bases, or very few, were detected at the end of the reads. No trimming was performed. Reads were aligned to reference genome *Rattus_norvergicus*. Rnor_6.0.87 using TOPHAPT and the alignment of the counts per read was performed using HTSeq*.* Differential gene expression analysis was performed using CUFFDIFF with replicates analysis to explore the differences in transcriptomic profiles between factor levels. All RNA-seq data sets generated and/or analysed during the current study were added to the Gene Expression Omnibus (GEO) under the accession number GSE158188.

Gene ontologies and pathways enrichment and overrepresentation were calculated with Metascape (11) (<http://metascape.org>) for every comparison and gene subset obtained. The Metascape analysis first identified all statistically enriched terms (can be GO/KEGG terms, canonical pathways, hallmark gene sets, etc., based on what your choice during the analysis), accumulative hypergeometric p-values and enrichment factors were calculated and used for filtering. Remaining significant terms were then hierarchically clustered into a tree based on kappa-statistical similarities among their gene memberships. Then 0.3 kappa score was applied as the threshold to cast the tree into term clusters. We then selected a subset of representative terms from this cluster and convert them into a network layout (see Figures 7 and 8). More specifically, each term is represented by a circle node, where its size is proportional to the number of input genes fall into that term, and its colour represents its cluster identity (i.e., nodes of the same colour belong to the same cluster). Terms with a similarity score > 0.3 are linked by an edge (the thickness of the edge is proportional to the similarity score). The network is graphically displayed with Cytoscape (v3.1.2) with “force-directed” layout and with edge bundled for clarity. One term from each cluster is selected to have its term description shown as label

**c-FOS immunohistochemistry after acute cocaine challenge**

When animals reached PND 90, they were injected either with cocaine (cocaine hydrochloride: 20 mg/kg i.p. Alcaliber, Spain.) or saline (1 mL/kg i.p. 0.9% NaCl sterile solution; Vitulia, Spain). Ninety minutes later they were anaesthetised with an injection of a 16% chloral hydrate solution (400 mg/kg i.p.) and transcardially perfused with PBS 0.1 M followed by 4% paraformaldehyde. Brains were then extracted and kept in fixing solution (4% paraformaldehyde) for 24 h and then transferred to a 30% sucrose solution for another 24 h. They were then kept at -20ºC in a glycerol/ethylene glycol (30%/30%) and (40%) PB 0.4M solution.

Fifty-microns coronal brain slices were obtained in a vibratome, transferred to a 30% glucose solution and kept at -4ºC for 24 h. They were then transferred to a -20ºC freezer and kept in a glycerol/ethylene glycol and PB 0.4 M solution until immunohistochemistry.

Free-floating tissue samples were washed in PBS 0.1 M (3 successive 10 min rinses) and then incubated with 0.3% hydrogen peroxide v/v in PBS at room temperature for 30 min. They were then incubated 1 hour in blocking solution (2% (v/v) normal goat serum + 0.3% (v/v) Triton-X 100 in PBST) and washed (PBST, 3x10 min). After this, they were incubated at 4°C for 24 hours in a rabbit c-Fos antibody (1:50000; Merck ABE457 Lot: 3116957). Sections underwent 3x10 min PBST rinses followed by a 1-hour incubation in biotinylated goat anti-rabbit IgG (1:200; Vectastain; BA-1000-1.5, LOT: ZE1218). After washing (PBS, 3x10 min), sections were incubated in ABC reagent (avidin-biotin complex kit, Vector Labs) at room temperature for 1 hour, washed (PBS, 3x10 min), and reacted in diaminobenzidine (DAB; approximately 5 minutes) to reveal neurons labelled for c-Fos in brown. Sections were then washed (PBS, 3x10 min). Sections were allowed to dry and mounted on microscope slides, and coverslipped using DPX.

Tissue images were captured at 10 X optical magnification using brightfield microscopy. The background was subtracted using a rolling ball procedure (radius 12.00), and the c-Fos positive cells per ROI were counted using the particle analysis option in ImageJ (size: 80-200, circularity: 0.50-1.00) by a researcher blind to the experimental conditions.

**Cocaine self-administration**

On PND 90, animals underwent a single food-reinforced FR1 instrumental training session limited to 10 reinforcers in Med-Associates operant boxes. After this, an intravenous polyvinylchloride tubing (0.064 mm i.d.) catheter was implanted into the right jugular vein at approximately the level of the atrium and passed subcutaneously to exit the mid scapular region(12). Surgical procedures were performed under isoflurane gas anaesthesia (5% for induction and 2% for maintenance) and buprenorphine analgesia (0.05 mg/kg s.c. Buprex, Indivior, U.K.). After surgery, the rats were allowed to recover for 8-10 days and a nonsteroidal anti-inflammatory drug (NSAID) (meloxicam - Metacam™: 15 drops of a 1.5 g/ml solution per 500 ml of water) was added to the drinking water during the first four days of recovery. Until the end of the self-administration procedure, the catheters were flushed daily with a sterile saline solution containing sodium heparin (100 IU/ml) and gentamicin (1 mg/ml) to maintain catheter patency and prevent infections.

The cocaine self-administration protocol was carried out in Coulbourn boxes (thus providing a different context to the one where the initial instrumental training took place). An active lever (AL) with different programmed contingencies and an inactive lever (IL) that always remained without programmed contingencies remained protracted during all the sessions. Over the AL there was a cue light that was turned on at the beginning of the session and turned off for 10 s (time out) at the beginning of an infusion (that lasted for 7 s), thus indicating cocaine availability. Cocaine (Alcaliber, Spain) infusions (0.5 mg/kg in 100 µL of sterile saline solution) were administered by an electronic pump (Harvard Apparatus, USA). Lever presses during time out (TOLP) did not have any programmed contingencies. The protocol consisted of 6 consecutive phases: (1) acquisition:12 daily sessions lasting 2 hours each under an FR1 schedule; (2) motivation for consumption: 6 sessions of 2 hours under a progressive ratio (PR) schedule(13); (3) reacquisition: 3 sessions of 2 hours under FR1; (4) compulsive (punished) seeking: a single 1-hour session under an FR3 schedule in which the animal randomly received an infusion or a 0.5 mA plant shock for 0.5 s. (5) extended access: 10 sessions of 6 hours each under FR1; (6) cue-induced relapse: 4 sessions of 1 hour each with cues as in the acquisition sessions but without drug delivery, occurring after 1, 30, 60 and 100 days of forced abstinence.

In the acquisition, reacquisition and extended access phases, main measurements were AL presses (ALP), TOLP, and IL presses (ILP). In motivation for consumption, we also registered the number of infusions achieved, the breaking point reached and a specific motivation index:

$$Motivation index =\frac{Infusions on 1st PR session}{Infusions on 1st PR session+Infusions on 12th AQ session}*100$$

In the compulsive consumption session, we also calculated a specific compulsivity index as:

$$Compulsivity index =\frac{\mathrm{Events}\left( shocks or infusions \right)}{Infusions in 1st hour of last RA session}*100$$

**SUPPLEMENTARY RESULTS**

**Bodyweight**

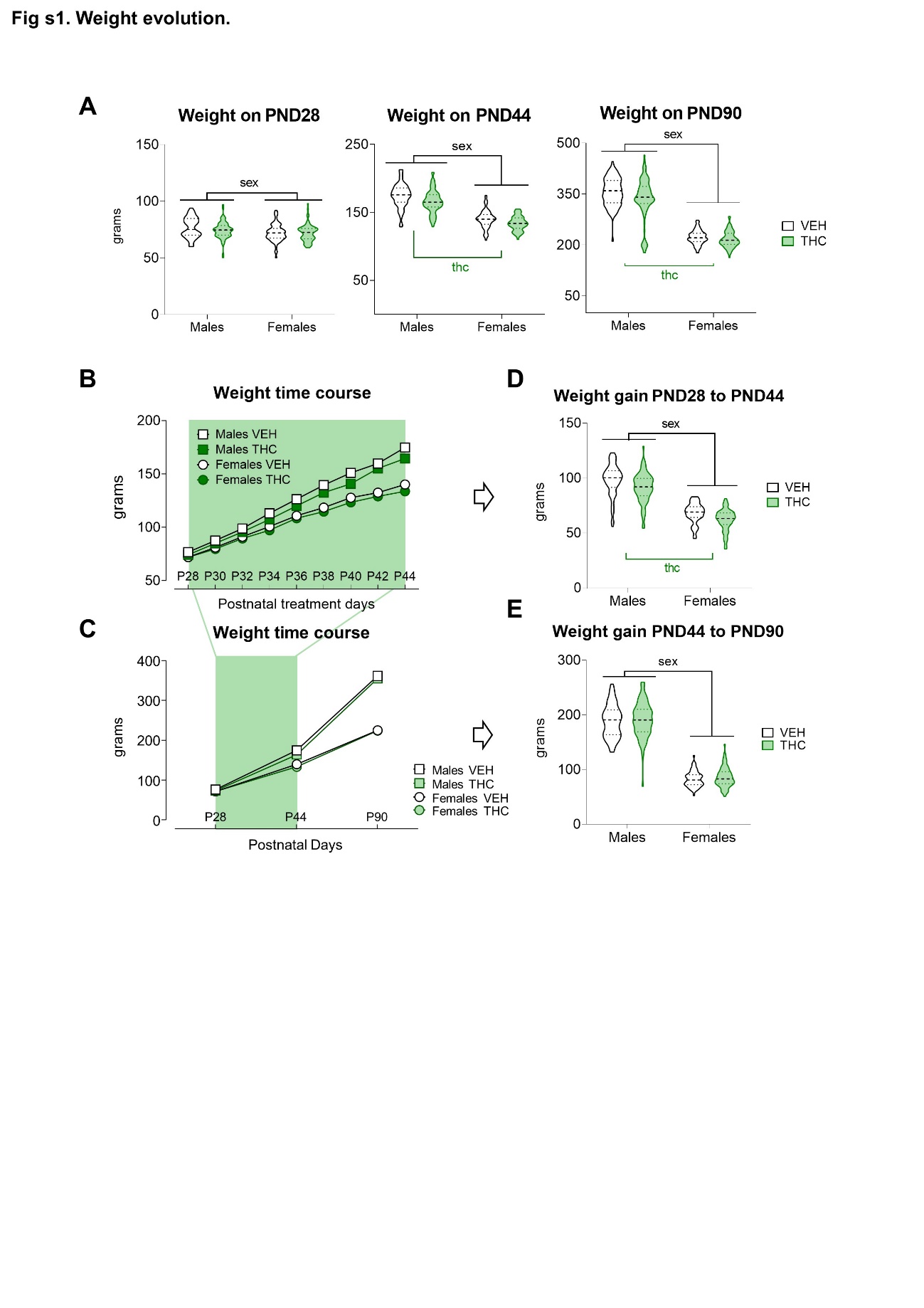
As expected, females had lower body weight than males at the beginning of the treatment (see Figure S1A). After the chronic treatment, THC-exposed rats (irrespective of the sex) had a marginally lower body weight than vehicle-exposed controls (Table S9). The increment in body weight was also lower during and after the treatment in females as compared to males and in THC-exposed animals as compared to vehicle controls (Figure S1B, D and Table S9).

**Figure S1. Bodyweight evolution.** Green coloured lines and “thc” represent a significant difference within levels due to the Adolescent Treatment factor. Black lines and “sex” represent significant difference within levels of the Sex factor. **A)** Bodyweight in three different time points. Only the expected sex differences were present in the first treatment day (PND28) (F_1,297_=14.686; p<0.000; η_p_^2^=0.04) indicating no prior weight bias due to Adolescent Treatment factor. Nonetheless, the mean body weight was lower THC groups at the end of the treatment (PND44) (F_1,299_=20.162; p<0.000; η_p_^2^=0.06) and at adulthood (PND90) (F_1,341_=14.686; p=0.004; η_p_^2^=0.02) **B)** Bodyweight time course across chronic treatment and **C)** Bodyweight time course in the three experimental points selected. **D)** Net body weight gain during treatment (PND44 Weight - PND28 Weight) and **E)** Net body weight gain during the washout period (Bodyweight on PND90– Bodyweight on PND44). The bodyweight gain during the THC treatment was slightly lower in the groups of THC-treated animals (F_1.296_=22.585; p<0.000; η_p_^2^=0.07). However, only a significant Sex effect (male>female) was found in weight gain during the washout period (F_1,294_=362.205; p<0.000; η_p_^2^=0.51)

**Pavlovian Conditioned Approach**

The repeated measures ANOVA showed a significant progression across the training session in the PCA index (see Figure S2 C, D). In the last training session, we observed a significant Treatment effect. This effect cannot be attributed to Treatment effects in the amount of CS^+^ LPs, ILPs, HE, CS^+^HE or MAG and CS^+^MAG on that session (see Table S2). Instead, it was mainly due to the effects of THC on the main variables that constitute the PCA index. Indeed, there was a significant Treatment effect in probability difference and latency score**,** and there was a trend towards statistical significance in the response bias**.** Another complementary variable was the first response probability which showed the same significant Treatment effect, whereby THC-exposed animals tended to first explore the magazine instead of interacting with the CS^+^ once the later appears (Figure S2D-G and Table S2).

This effect was also significant along with the training sessions, but the components of the PCA index did not show any Treatment effect in the repeated measures analysis. With regard to sex differences, females, independently of the Adolescent Treatment, spent more time seeking the reinforcer in the food magazine than their counterparts as reflected by the Sex effect on CS+HE, MAG and CS+MAG in the cumulative data across the training sessions (Table S2).

The classification of animals into goal-trackers or sign trackers also revealed, as expected, that more animals fell into the first category in the groups exposed to THC during adolescence (Figure S4B). The control groups had a majority of animals classified as intermediate.

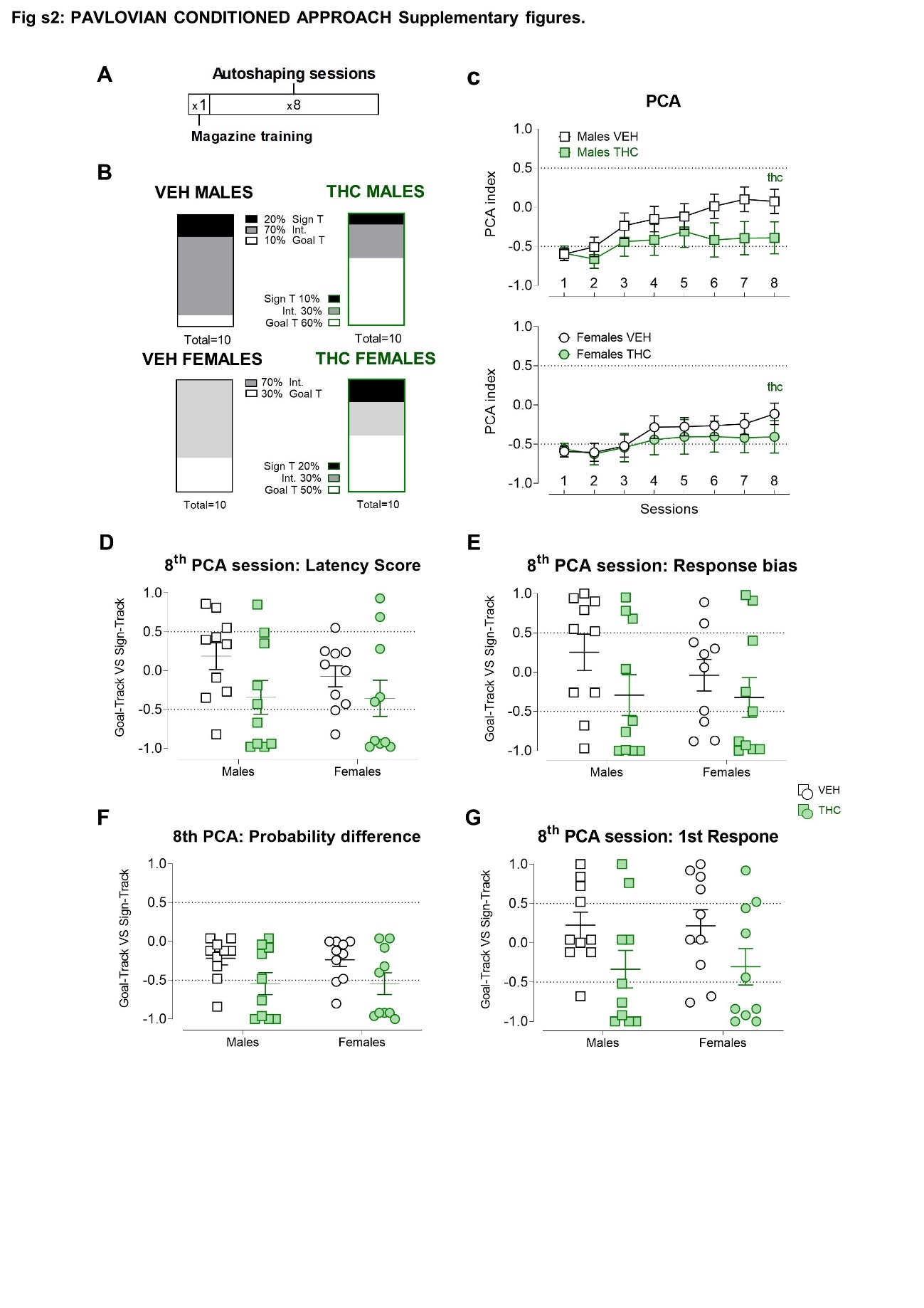

**Figure S2:** **Pavlovian Conditioned Approach Supplementary figures.**

**A)** Timeline of experimental phases. **B)** Percentual distribution of the three different PCA clusters in each group. **C)** PCA index across the eight auto-shaping sessions. **D), E)** and **F)** Individual components calculated within each session to derive the global PCA index. Graphs show the distribution in each component in the 8^th^ session. An increase in goal- tracking behaviour in THC animals was significant in the Probability difference (F_1,36_=7.397 ; p=0.010; η_p_^2^=0.17 ) and Latency score (F_1,36_= ; p= 0.043; η_p_^2^=0.11 ) indices but did not reach statistical significance in Response bias (F_1,36_=3.058; p=0.089; η_p_^2^=0.08). **G)** Another additional proxy, not included in the general index was also calculated, namely the First response, that also showed a to be significantly affected in the same THC rats (F_1,36_= 6.447; p= 0.016; η_p_^2^= 0.15).

**Pavlovian-to-Instrumental Transfer**

Across Pavlovian training sessions, CS+HEs ratio showed a significant Sex x Adolescent Treatment interaction, but there were no significant simple effects (see Table S3). All groups progressively increased their bias to perform more HEs during CS^+^ than during CS^-^ and finished the Pavlovian training with ratios over 0.7 (see Figure S3B). Females seemed to have a higher locomotor activity as they made more HEs (both during CS^+^ and CS^-^) during Pavlovian training (see Table S3). There was also a trend for females to make more HEs during the ISI (see Table S3).

Along the time course of the instrumental training sessions, animals rapidly learned instrumental contingencies associated with both levers. The lever press ratio at the end of the training was over 0.8 (see Figure S3C). Females took more time to achieve the limit of reinforcers each session and performed less TO ALPs during the instrumental training (Table S3). No effect in the number of head entries across the sessions was found.

There was a decrease in the number of ALP across the extinction sessions. The repeated measures ANOVA showed a Sex x Adolescent Treatment interaction on ILPs, but no significant simple effects were present (Table S3).

During the PIT testing session, the two-way ANOVA of the subjects that expressed a CS^+^ ALP ratio over 0.5, resulted in a global effect of THC and a Sex x Adolescent Treatment interaction (Table S3). The simple effect analysis showed that THC-exposed males had a higher CS^+^ ALP ratio compared with their controls. This effect mainly resulted from a lower CS^-^ ALPs in the group of THC-exposed males without differences in the rate of CS^+^ ALPs as a result of THC (see Figure S3D, E and F). The CS^+^ HE ratio also showed a significant Sex x Adolescent Treatment interaction resulting from the increased ratio in THC females, as compared to the VEH females (Figure S3G, H and I).

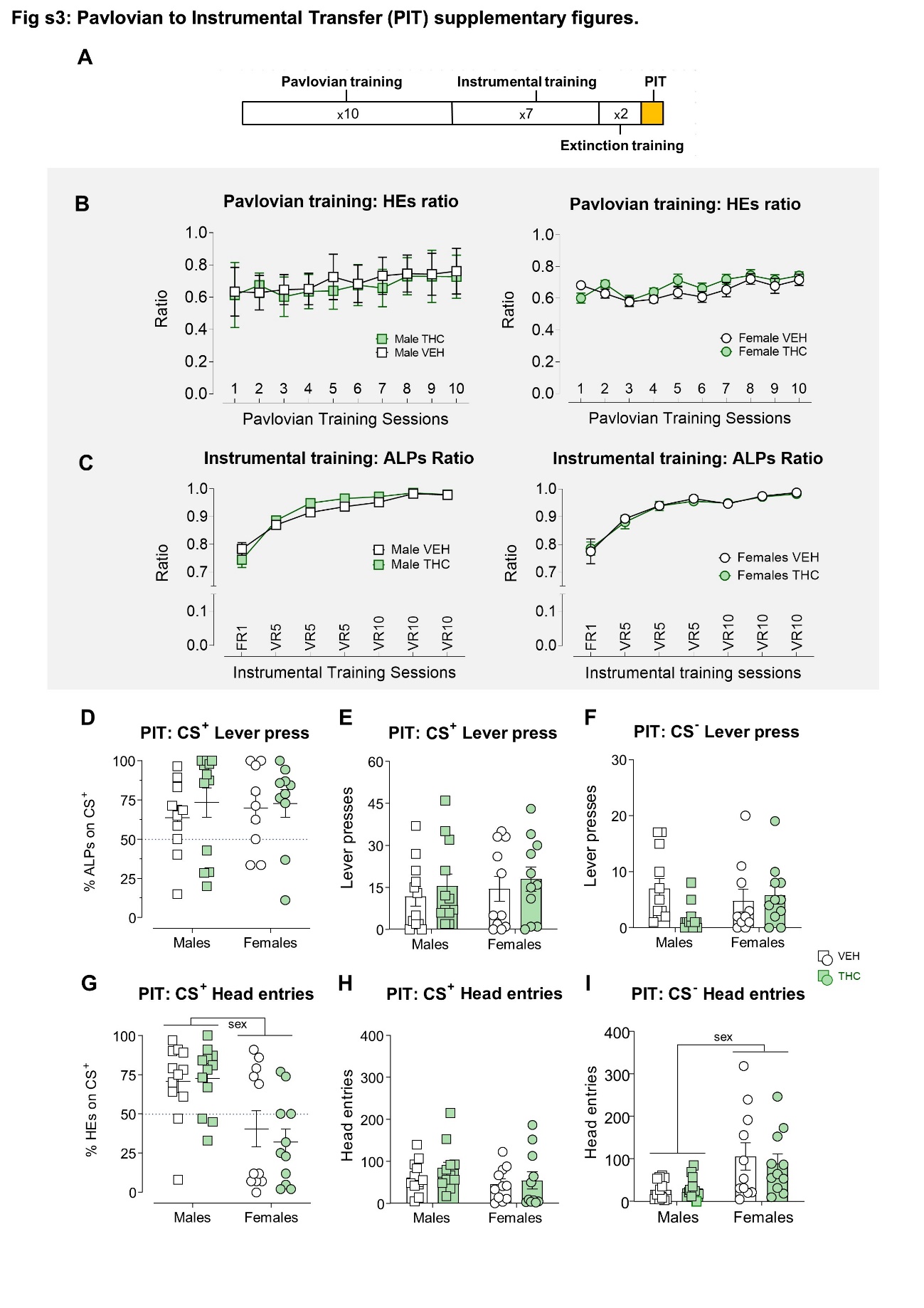

**Figure S3. Pavlovian to Instrumental Transfer (PIT) supplementary figures.**

**A)** Timeline of experimental phases. **B)** Head entries (HEs) ratio across pavlovian training sessions. Males and females are plotted separately for the sake of visual clarity. There was a general increase in the amount of HEs during CS+ across the sessions (F_5.7,341.6_=9.657; p=0.000; η_p_^2^=0.14). Also, there was a Sex x Adolescent Treatment interaction (F_5.7,341.6_ =2.338; p=0.033; η_p_^2^=0.03), but no differences were detected in the analysis of the simple effects. **C)** Active to inactive Lever Press ratio across training sessions. Males and females plotted separately for the sake of visual clarity. There was a general increase in the discrimination and preference for the AL over the IL across the sessions (F_2.09,396_=136.664; p=0.000; η_p_^2^=0.67). **D)** Percentage of lever presses during the CS+ in the transference test. No differences between groups were detected, including animals that did not express PIT (F_3,43_=1.031; p=0.389; η_p_^2^=0.07). **E)** Total Lever Presses during the CS+ in the transference test. No differences between groups were detected (F_3,42_=0.385; p=0.764; η_p_^2^=0.02). **F)** Total Lever Presses during the CS- in the transference test. **G)** Percentage of HEs during the CS+ in the transference test. Females made relatively fewer HEs during the CS+ as compared to the CS- than males (F_1,45_=18.095; p=0.000; η_p_^2^=0.301). The mean percentage of HEs during the CS+ was clearly above 50% in the males, thus indicating clear conditioning of the CS+ while in the females, this value was below 50%. **H)** Total HEs during the CS+ in the transference test. No differences between groups were detected (F_3,42_ =1.085; p=0.366; η_p_^2^=0.07). **I)** Total head entries during the CS- in the transference test. Females made more HEs during the CS- presentations than the males (F_1,42_=12.460; p=0.001; η_p_^2^=0.22).

**Habit Formation**

During the short training instrumental training sessions, animal successfully acquired lever-pressing behaviour as suggested by the significant effect of the Sessions factor (see Table S5). However, there were no significant effects of Sex or Adolescent Treatment. Additionally, we found no effects along with short training sessions as a result of Sex or Adolescent Treatment factors in any of the measurements analysed. After short training, females performed more lever presses in non-devalued conditions, and THC exposure also increased the rate of responding in these animals. Indeed, the simple effects analysis showed a significant increment of lever presses in THC-females compared with their controls and also with their male counterparts. A similar pattern of results was evident in the devalued condition. HEs also showed a trend to be incremented in THC animals again with this increased activity mainly found in THC-females (see Table S5).

Across the extended training sessions, we only found a Sex x Adolescent Treatment x Sessions interaction in the HEs, pointing to a sex effect in THC animals whereby there fewer HEs performed by THC-females as compared to THC-males. There were no effects in the difference in lever presses in the devalued vs non-devalued conditions, but the time spent in the magazine time showed a significant Sex x Adolescent Treatment interaction whereby THC-males spent more time inspecting the magazine than VEH-males, and this effect was absent in the females. Moreover, there was a Treatment effect in the time spent in the magazine during the omission training; THC-exposed animals spent more time interacting with the food magazine (an effect reminiscent of the increased goal-tracking behaviour shown in the PCA experiment; see Figure 1A).

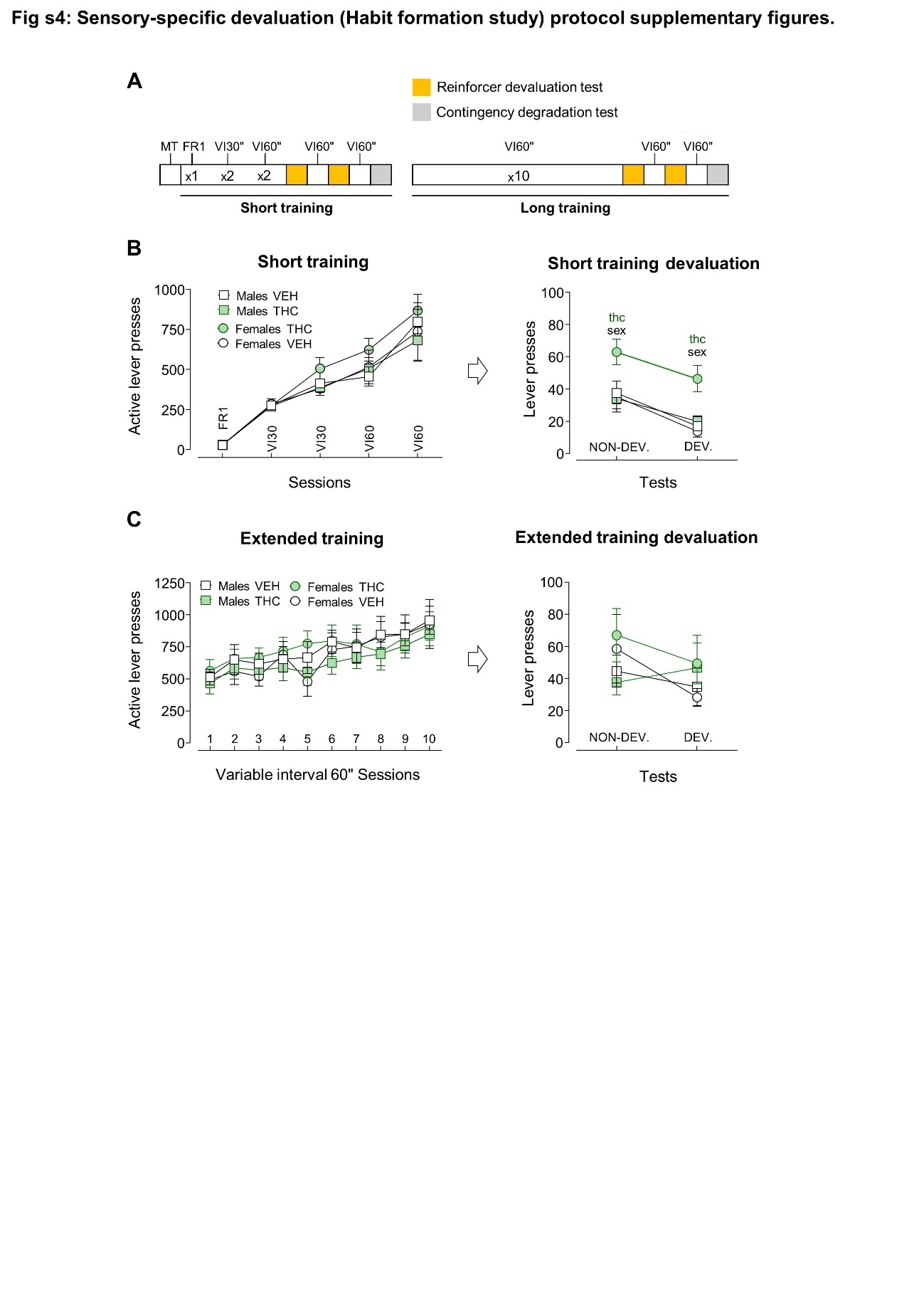

**Figure S4: Habit formation study protocol supplementary figure**.

**A)** Timeline of experimental phases. **B)** Lever presses during short training sessions and sensory-specific satiety outcome devaluation test. No effect of Sex or Adolescent Treatment were observed across training sessions. In the test sessions, there was an increased lever pressing in THC females in both conditions compared with VEH females (F_1,18_= 10.740; p= 0.004; η_p_^2^= 0.37) and THC males (F_1,18_= 9.526; p= 0.006; η_p_^2^= 0.35). All the animals decreased their responding in the devalued condition (suggestive of goal-directed behaviour and the absence of habitual responding) (F_1,36_= 30.976; p<0.000; η_p_^2^= 0.37) **C)** Extended training sessions and sensory-specific satiety outcome devaluation test. All groups progressively increased their responding across the training sessions (F_4.48,147.72_ = 21.575; p<0.000; η_p_^2^= 0.39). There were no session effects in LP in the tests, indicating the absence of devaluation and probably the development of a stimulus-response guided behaviour compatible with a habit. There were no Sex or Adolescent Treatment effects (F_1,35_ = 1.294; p=0.263; η_p_^2^= 0.03).

**2-Choice Serial Reaction Time Task**

The number of training sessions necessary to achieve a stable baseline showed a trend of the Treatment to have a significant effect (0.055), suggesting that THC-exposed animals required fewer training sessions to reach this a stable baseline (Table S5; Figure S5B).

During the six baseline sessions, before long ITI sessions, there was a lower CR rate in THC animals; IR showed a specular trend, that is, an increased IR in THC-treated animals. OR, PerR, TOR and HE were not different across the groups (see Table S4).

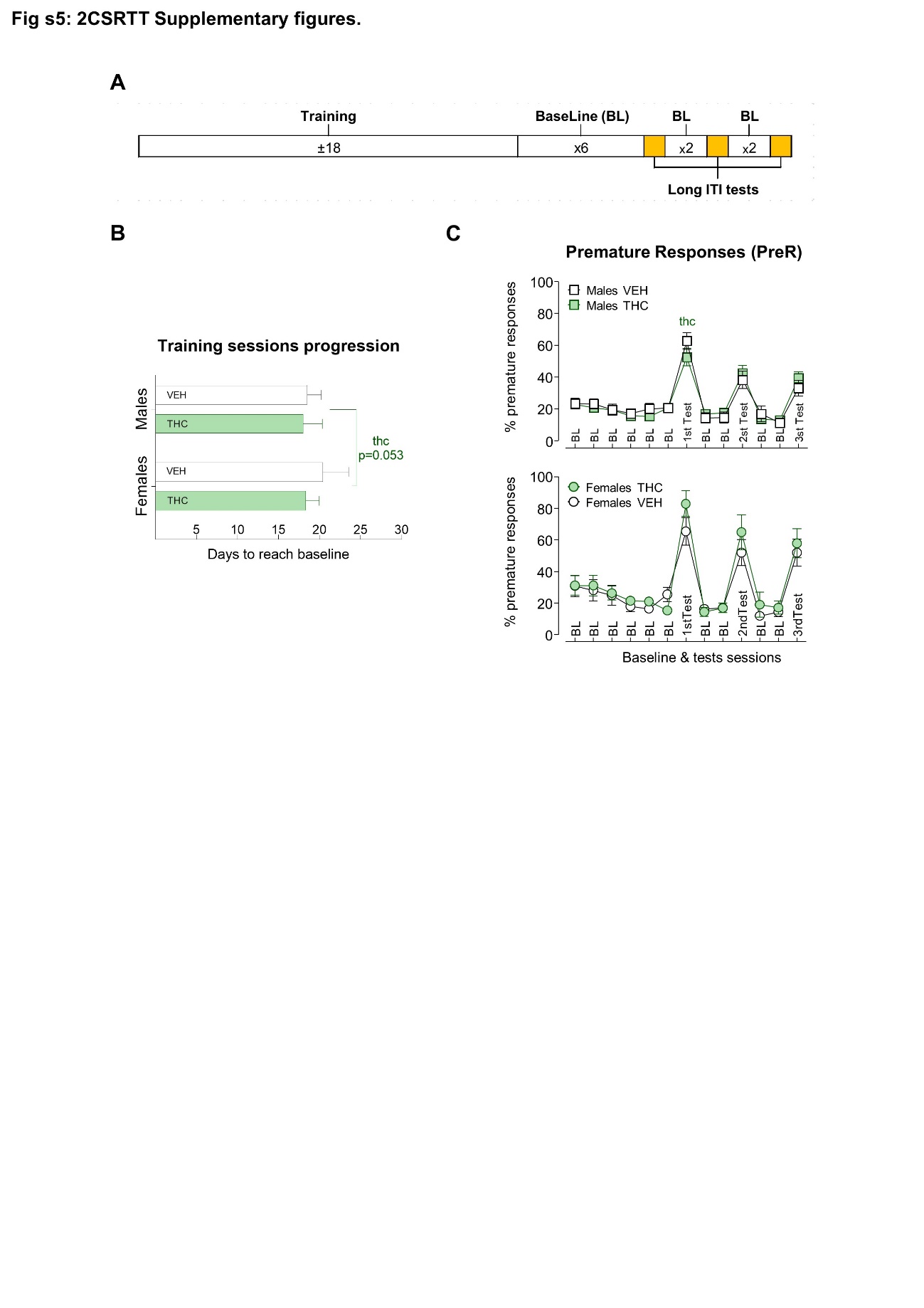
A preliminary analysis of the crude premature responses in the longITI sessions, revealed an effect of the Treatment that disappeared in the subsequent testing sessions, whereby THC-males showed fewer PreR than VEH-males. However, when we analysed the last three training sessions in order to verify that the number of PreR was stable enough to provide a valid baseline, we found a significant Sessions x Treatment interaction (see Figure S5C and Table S4), so we decided to calculate the percentage of change in the longITI sessions as compared to the precedent baseline sessions (see Figure 1E and the supplementary methods in this file).

**Figure S5: 2CSRTT Supplementary figure.**

**A)** Timeline of the experimental phases. **B)** Number of training days required to reach the baseline sessions. THC animals showed a trend (F_1,44_=3.888; p=0.055; η_p_^2^=0.08) to require fewer days to reach baseline sessions than VEH rats. No significant effect of Sex was detected (F_1,44_=2.949; p=0.093; η_p_^2^=0.06). **C)** Percentage of premature responses in the final experimental stages. This includes the six baseline sessions, the three long-ITI tests and the re-baseline sessions in between them. In the first long-ITI test, a Sex x Adolescent Treatment interaction (F_1,44_=7.483; p=0.009; η_p_^2^=0.14) and further simple effects analysis showed an effect of sex due to an increase in PreR in females VEH compared to males VEH (F_1,44_=7.630; p=0.008; η_p_^2^=0.14) and a decrease in PreR in Male THC compared to males VEH (F_1,44_=5.740; p=0.021; η_p_^2^=0.11).

**cFOS Immunohistochemistry**

In addition to the results reported in the main body of the article, there was a significantly higher accumulation of c-Fos after cocaine (Adult Treatment effect) in the lateral orbitofrontal, cingulate, motor, insular and entorhinal cortices (see Table S6). This effect was also observed in the amygdala. The effects of cocaine were restricted to the males in the lateral orbitofrontal cortex. There were some interactive effects in the medial septal nucleus between the sex of the animal and cocaine/saline exposure but the simple effects did not reach statistical significance or in the Islands of Calleja where cocaine induced the expression of the c-Fos protein only in the females (See Figure S6 and Table S6). We have also detected THC-related effects in retrosplenial, somatosensory and piriform cortices where THC animals had higher levels of c-Fos proteins as compared to animals exposed to VEH. In the last two cases, this effect was only observed in the males (significant Sex x Adolescent Treatment interactions with significant simple effects in the males). We have also observed general main effects of the sex of the animals in the medial orbital, prelimbic, cingulate, motor, insular, entorhinal and retrosplenial cortices and in the Islands of Calleja with females showing higher values than males (see Table S6 and Figure S6).

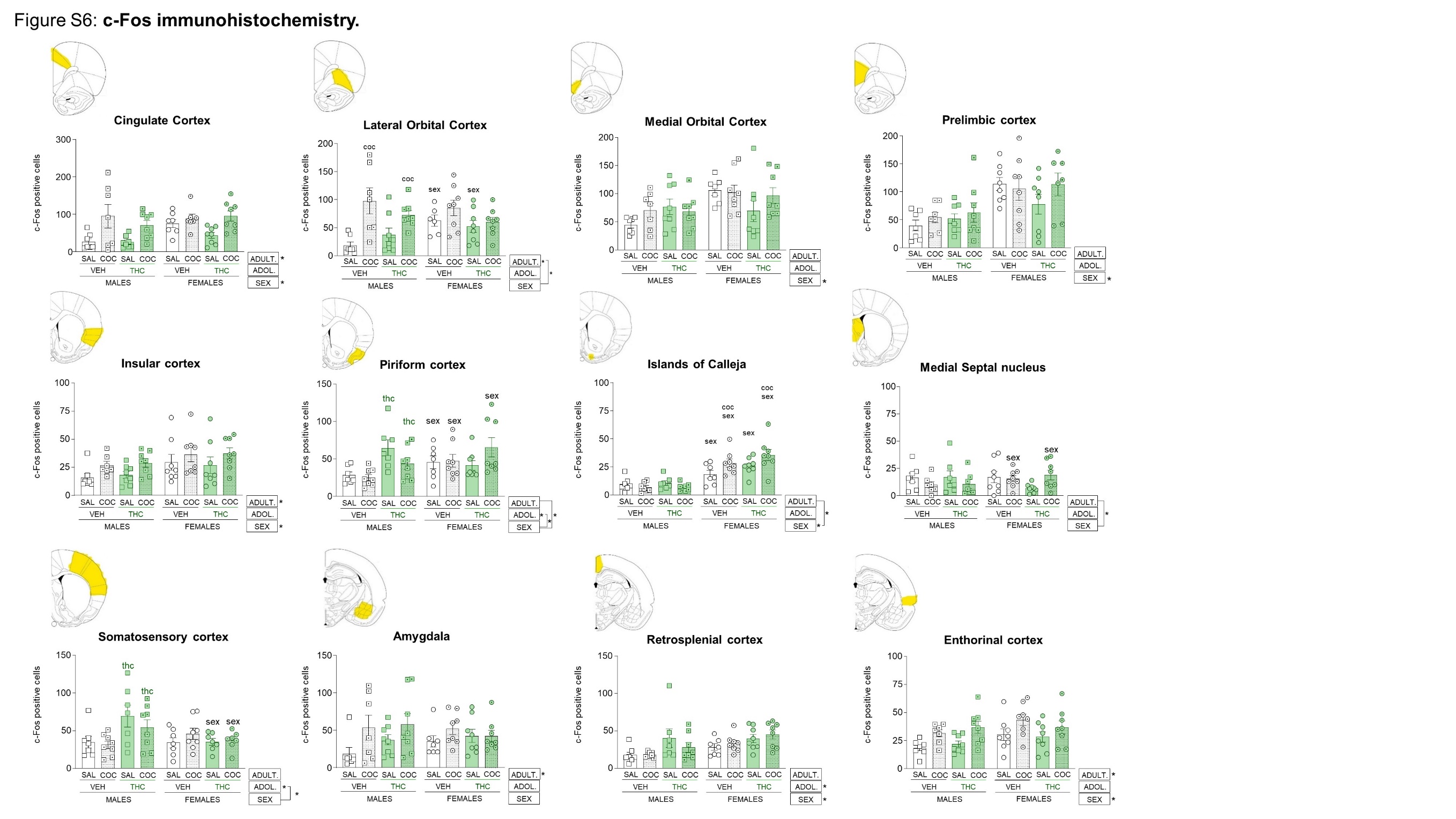

**Figure S6: c-Fos immunohistochemistry.** Brain areas showing significant effects Sex, Adolescent Treatment or Adult Treatment (cocaine injection) or interactions among them. Factor effects are indicated with “ * ” next to the legend boxes on the bottom right; interactions between factors are indicated with lines joining the factors and “ * ”. Significant results of the analysis of the simple effects of the interactions are indicated with “sex”, “thc” or “coc” on top of the experimental group. “sex” stands for differences with the corresponding group of the other sex, “thc” indicates a difference with the corresponding group of the other adolescent treatment (VEH) and “coc” indicates a significant difference with the saline group within the same adolescent treatment group.

**Cocaine Addiction-like Behavioural Screening**

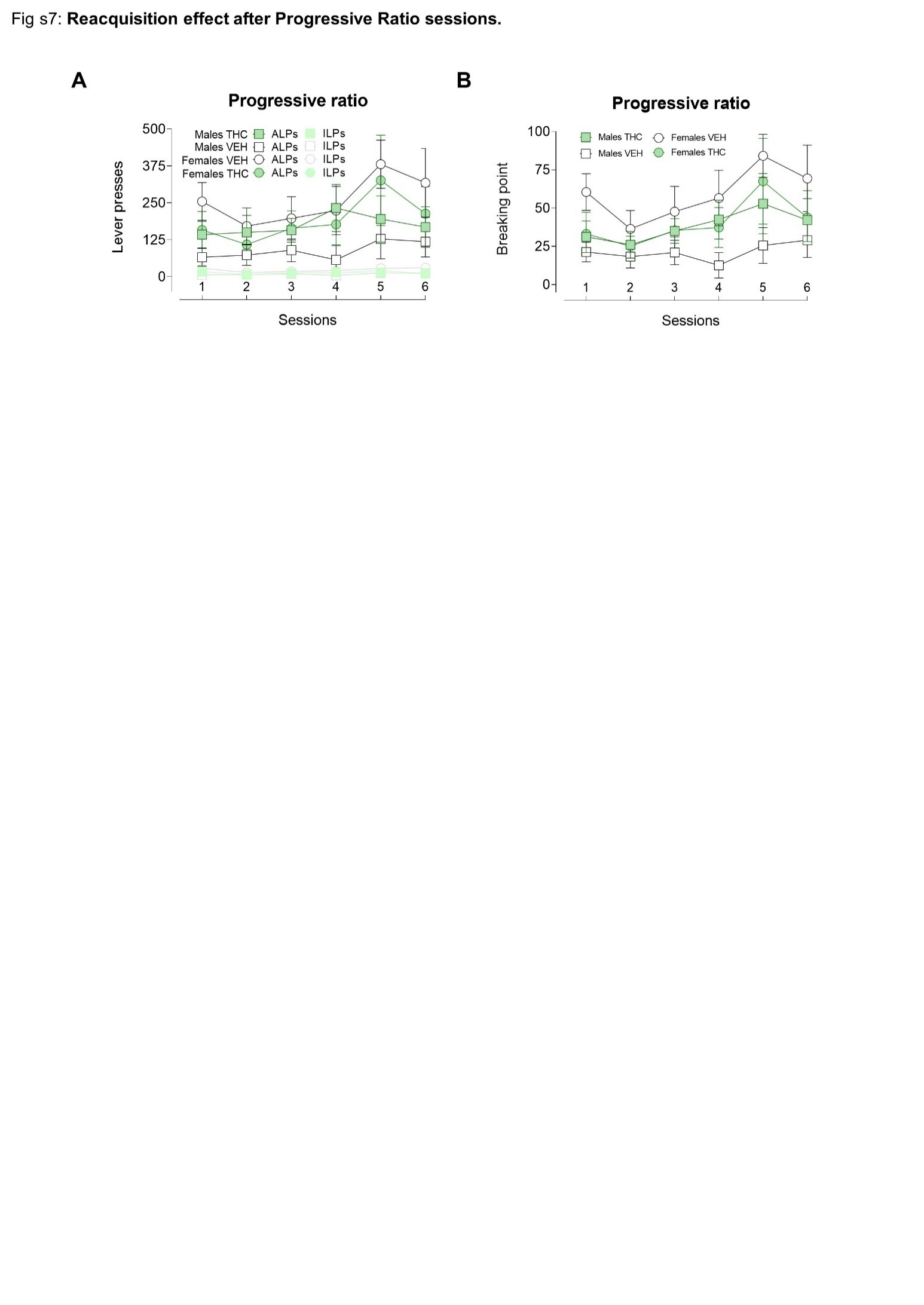
*Progressive ratio*: In the first day of progressive ratio (PR) schedule, there was a significant Sex x Adolescent Treatment interaction which suggested that males exposed to THC earned a higher number of infusions than their VEH controls whereas the opposite pattern emerged in the females. However, the follow up of this interaction did not show statistically significant effects (see Table S7). The analysis of the ALPs or the BPs across the six PR sessions only showed a significant effect of the Sessions factor (Figure S7A, B and Table S7.

**Figure S7: Cocaine self-administration progressive ratio performance.**

Mean values are depicted with circles or squares joined by lines in the repeated measures graphs. Error lines express the SEM. **A)** Lever presses during progressive ratio (PR) sessions. There was a progressive increase in the ALPs across the six PR sessions (F_5,145_ = 3.857; p=0.003; η_p_^2^= 0.11) that was similar across the groups. **B)** Breaking points achieved across the six PR sessions. There was a progression in the breaking points achieved across sessions (F_2.77, 80.28_=4.763; p=0.005; η_p_^2^=0.14).

**REFERENCES**

1. Frye GD, Breese GR. An evaluation of the locomotor stimulating action of ethanol in rats and mice [Internet]. *Psychopharmacology (Berl).* 1981;75(4):372–379.

2. Reagan-Shaw S, Nihal M, Ahmad N. Dose translation from animal to human studies revisited. *FASEB J.* 2008;22(3):659–661.

3. Ridgeway G, Kilmer B. Bayesian inference for the distribution of grams of marijuana in a joint [Internet]. *Drug Alcohol Depend.* 2016;165:175–180.

4. Chandra S et al. New trends in cannabis potency in USA and Europe during the last decade (2008–2017) [Internet]. *Eur. Arch. Psychiatry Clin. Neurosci.* 2019;269(1):5–15.

5. Holmes NM, Marchand AR, Coutureau E. Pavlovian to instrumental transfer: a neurobehavioural perspective. [Internet]. *Neurosci. Biobehav. Rev.* 2010;34(8):1277–95.

6. Bari A, Dalley JW, Robbins TW. The application of the 5-choice serial reaction time task for the assessment of visual attentional processes and impulse control in rats. [Internet]. *Nat. Protoc.* 2008;3(5):759–67.

7. Paxinos G, Watson C. The rat brain in stereotaxic coordinates (6th ed.). *Acad. Press* 2007;

8. Pascau J et al. Automated method for small-animal PET image registration with intrinsic validation [Internet]. *Mol. Imaging Biol.* 2009;11(2):107–113.

9. Paxinos G, Watson C. The Rat Brain in Stereotaxic Coordinates. 6th Edition2007;

10. Kallio MA et al. Chipster: User-friendly analysis software for microarray and other high-throughput data. *BMC Genomics* 2011;12(507). doi:10.1186/1471-2164-12-507

11. Zhou Y et al. Metascape provides a biologist-oriented resource for the analysis of systems-level datasets. *Nat. Commun.* 2019;10(1):1523.

12. Higuera-Matas A et al. Augmented acquisition of cocaine self-administration and altered brain glucose metabolism in adult female but not male rats exposed to a cannabinoid agonist during adolescence. *Neuropsychopharmacology* 2008;33(4). doi:10.1038/sj.npp.1301467

13. Sánchez-Cardoso P et al. Modulation of the endogenous opioid system after morphine self-administration and during its extinction: a study in Lewis and Fischer 344 rats. [Internet]. *Neuropharmacology* 2007;52(3):931–948.
